# Architecture of the human erythrocyte ankyrin-1 complex

**DOI:** 10.1101/2022.02.10.479914

**Authors:** F. Vallese, K. Kim, L.Y. Yen, J.D. Johnston, A.J. Noble, T. Calì, O.B. Clarke

## Abstract

The stability and shape of the erythrocyte membrane is provided by the ankyrin-1 complex, but how it tethers the spectrin-actin cytoskeleton to the lipid bilayer and the nature of its association with the band 3 anion exchanger and the Rhesus glycoproteins remains unknown. Here we present structures of ankyrin-1 complexes purified from human erythrocytes. We reveal the architecture of a core complex of ankyrin-1, the Rhesus proteins RhAG and RhCE, the band 3 anion exchanger, protein 4.2 and glycophorin A. The distinct T-shaped conformation of membrane-bound ankyrin-1 facilitates recognition of RhCE and unexpectedly, the water channel aquaporin-1. Together, our results uncover the molecular details of ankyrin-1 association with the erythrocyte membrane, and illustrate the mechanism of ankyrin-mediated membrane protein clustering.

## INTRODUCTION

An ordered lattice of spectrin and actin gives mechanical stability and shape to otherwise fragile cellular membranes. The periodic spectrin-actin cytoskeleton was first identified in erythrocytes^1^, but has since been shown to underlie the morphology and subcellular organization of other cell types, including neurons^2^. The spectrin-actin network is tethered to the membrane by giant, spring-like proteins known as ankyrins, which also serve the purpose of clustering diverse membrane proteins at specific sub-cellular locations.

Three ankyrin proteins are present in vertebrates: ankyrin-1 (ankyrin-R), ankyrin-2 (ankyrin-B) and ankyrin-3 (ankyrin-G). Ankyrins share a common domain arrangement, consisting of a highly conserved N-terminal membrane binding domain with 24 ankyrin repeats (AR1-24), followed by a spectrin binding module (ZU5-ZU5-UPA; ZZU) and a death domain of unclear functional significance. The characteristic 24-repeat membrane binding domain of ankyrins has a groove on the inner concave surface that mediates recruitment of linear peptide motifs from target proteins, as well as mediating autoinhibition by regions of the MBD-ZZU linker and the C-terminal tail. Ankyrin-1 was first identified in erythrocytes^3^, where it mediates the attachment of the band 3 anion transport protein (AE1; SLC4A1) to spectrin^4^. It is also found in other tissues, including skeletal muscle and brain, where it has recently been shown to mediate clustering of Kv3.1b K+ channels in specific neuronal subtypes^5^. Ankyrin 2 and 3 play pivotal roles as dynamic adaptor platforms in cardiomyocytes and striated muscle cells^6^, and in neurons of the central^7^ and peripheral^8^ nervous systems, respectively.

Conserved, linear ankyrin binding motifs have been identified in multiple classes of ankyrin target proteins, including voltage gated sodium (NaV) channels^9^, Na+/K+ ATPases^10^, and the band 3 anion exchanger^11^. These motifs are believed to mediate binding of targets via interaction with the inner groove on the concave face of ankyrin^12^. There is evidence that ankyrin can bind multiple targets at once^13^, but the mechanism by which an ankyrin molecule with a single peptide binding groove can recruit multiple targets simultaneously remains unclear – do multiple peptides bind in different regions of the groove at the same time, and how does this comport with the capacity of ankyrins to recruit multiple copies of the same protein?

In erythrocytes, ankyrin-1 attaches the cytoskeleton to the membrane via direct interaction with spectrin^14^, and additionally mediates clustering of integral membrane proteins, helping maintain the structure of the membrane and coordinating the spatial organization of transporters and channels involved in the regulation of cellular volume and cytosolic composition. Mutations in components of the erythrocyte ankyrin-1 complex are linked to an inherited anemia known as hereditary spherocytosis (HS), in which the characteristic biconcave disc shape of healthy erythrocytes is lost, resulting in small, fragile, spherical cells known as spherocytes^15^. Ankyrin and spectrin mutations are also associated with several other genetic diseases, including neurological and cardiac abnormalities^16^. The erythrocyte ankyrin-1 complex is known to contain the band 3 chloride/bicarbonate exchanger, the adaptor molecule protein 4.2, the single transmembrane protein glycophorin A (GPA), and the Rhesus proteins RhAG and RhCE^17^. However, the stoichiometry and architecture of the complex remains unknown, making it close to impossible to reconstitute recombinantly.

Here, we purified the ankyrin-1 complex from human erythrocytes, and captured a series of high- resolution structures by single particle cryogenic electron microscopy (cryo-EM). These revealed its architecture and composition, shedding light on the mechanism of membrane protein recruitment by ankyrin and protein 4.2, and identifying aquaporin-1 as an unanticipated component of the ankyrin-1 complex.

## RESULTS

### Purification and structure determination of the ankyrin-1 complex

The ankyrin-1 complex was purified from digitonin-solubilized human erythrocyte ghost membranes by density gradient centrifugation, followed by size exclusion chromatography (**Fig. 1a** and **Extended Data Fig. 1a-b**). The final sample contained a major 1.2 MDa species as assessed by mass photometry (**Extended Data Fig. 1c**). The presence of major components of the erythrocyte ankyrin-1 complex, including ankyrin-1, band 3, protein 4.2, glycophorin A and RhAG was assessed by SDS-PAGE and confirmed by mass spectrometry analysis (**Extended Data Fig. 1c**).

**Fig. 1:**
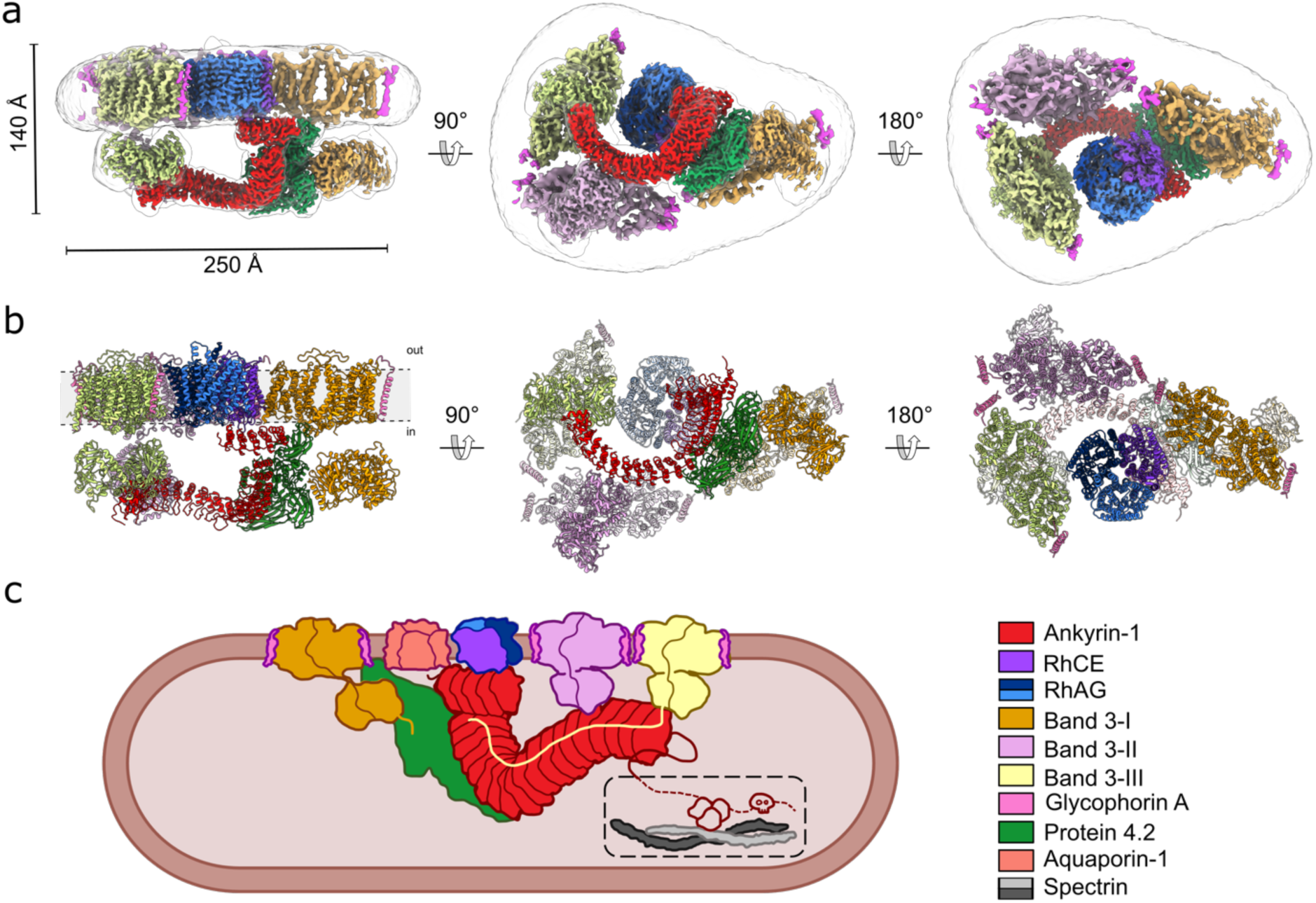
Architecture of the human erythrocyte ankyrin-1 complex. **a,** The overall cryo-EM density map of the ankyrin-1 complex (Class 1; transparent white surface) shown in three different views, with local reconstructions of different regions superposed. Band 3-I, II, III are colored orange, lilac and yellow respectively; ankyrin is colored red, protein 4.2 green, RhCE purple and RhAG blue and light blue. **b,** Model of the ankyrin-1 complex in the plane of the membrane (left panel), viewed from the cytoplasm (central panel) and from the outside of the plasma membrane (right panel). Reference colors are the same used in panel A. **c,** Schematic of the proposed ankyrin- 1 complex in the RBC. Inside the dashed line the connection between ankyrin and spectrin is depicted.

Cryo-EM analysis of the purified sample revealed both a mixture of ankyrin-1 containing complexes, as well as smaller complexes of free band 3 with glycophorin A (**Extended Data Fig. 2a**).

**Fig. 2:**
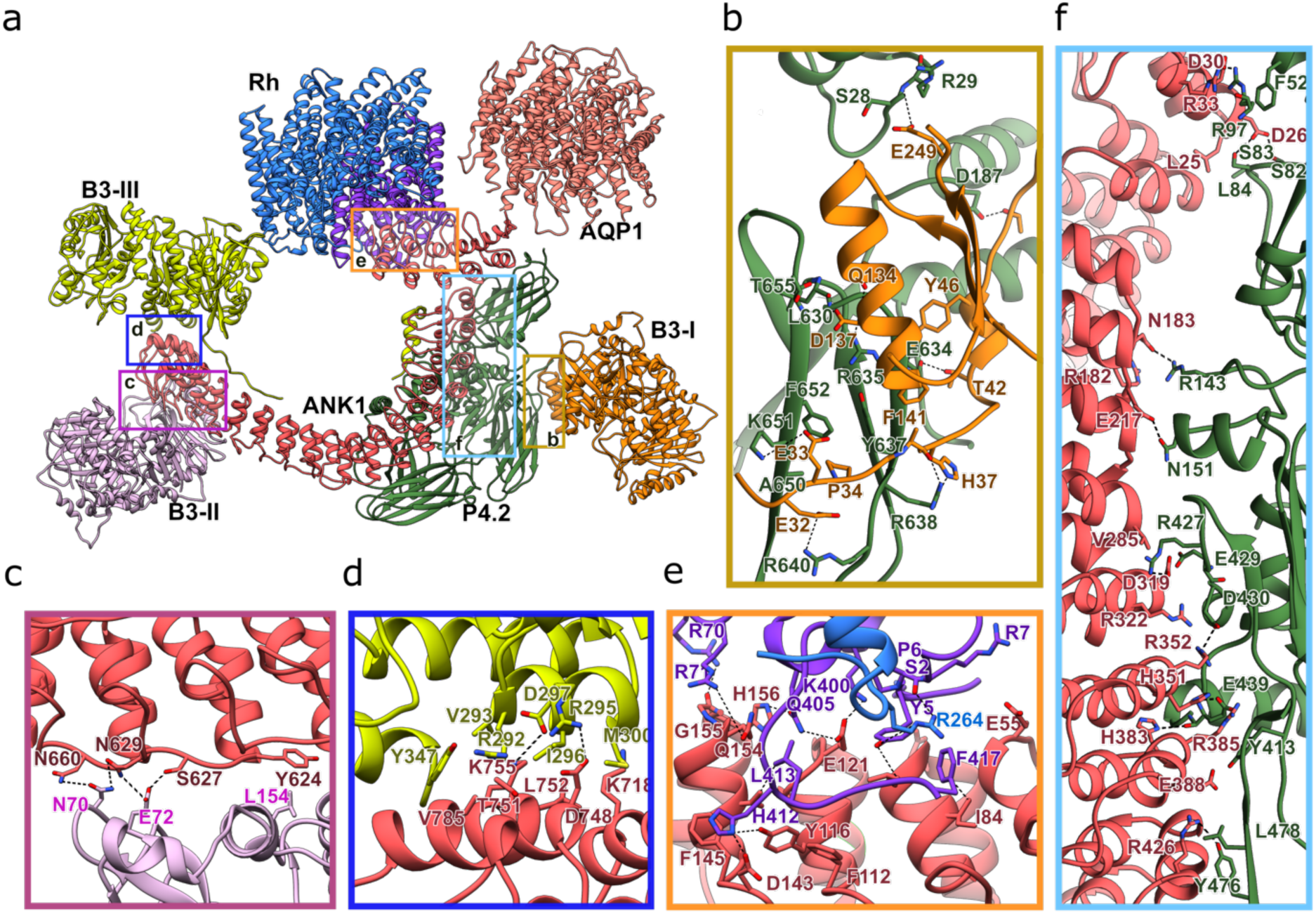
Interactions between ankyrin-1 and the proteins in the ankyrin-1 complex. **a,** Overview of ankyrin-1 interactions with binding partners in the ankyrin-1 complex. The proteins are depicted in cartoon representation. Transmembrane domains of band 3 and GPA are omitted for clarity. Ankyrin (red) interacts with RhCE (purple) and AQP1 (salmon), and both interact with the peptide binding groove of AR1-5; protein 4.2 (green) interacts with the exterior surface of AR1 and AR6-13; Band 3-II (lilac) and Band 3-III (yellow) interact predominantly with the exterior surface of AR17-24, N-terminus domain of Band 3-III interacts also with the peptide binding groove of AR6-10. Insets refer to regions highlighted in panels B to F. **b-f,** Close-up views of the interactions between protein 4.2 and Band 3-I (orange) (**b**); ankyrin and Band 3-II (**c**); ankyrin and Band 3-III (**d**); ankyrin and RhCE (**e**) and ankyrin and protein 4.2 (**f**). The key residues that mediate the interactions in the interfaces are shown as sticks. Key interactions are indicated with black dotted lines.

In addition to the major classes corresponding to higher molecular weight complexes with ankyrin, we also observed a significant class of particles corresponding to free band 3 dimers with high resolution features evident in the 2D classes (**Extended Data Fig. 2b**). *Ab initio* reconstruction and refinement of this class gave a C1 reconstruction at 3.2 Å with asymmetrically disposed cytoplasmic domains, similar to what we observed in the intact complex. Inspection of the map revealed that both protomers were in the outward open state, and that the transmembrane domain conforms to apparent C2 symmetry. Refinement of all band 3 glycophorin A dimers gave a 2.8 Å reconstruction with C2 symmetry, allowing us to build a complete atomic model of the dimeric transmembrane domain of band 3 in the outward open state, with each protomer of band 3 bound to one glycophorin A monomer (**Extended Data Fig. 2c, Extended Data Fig. 3-4**).

**Fig. 3:**
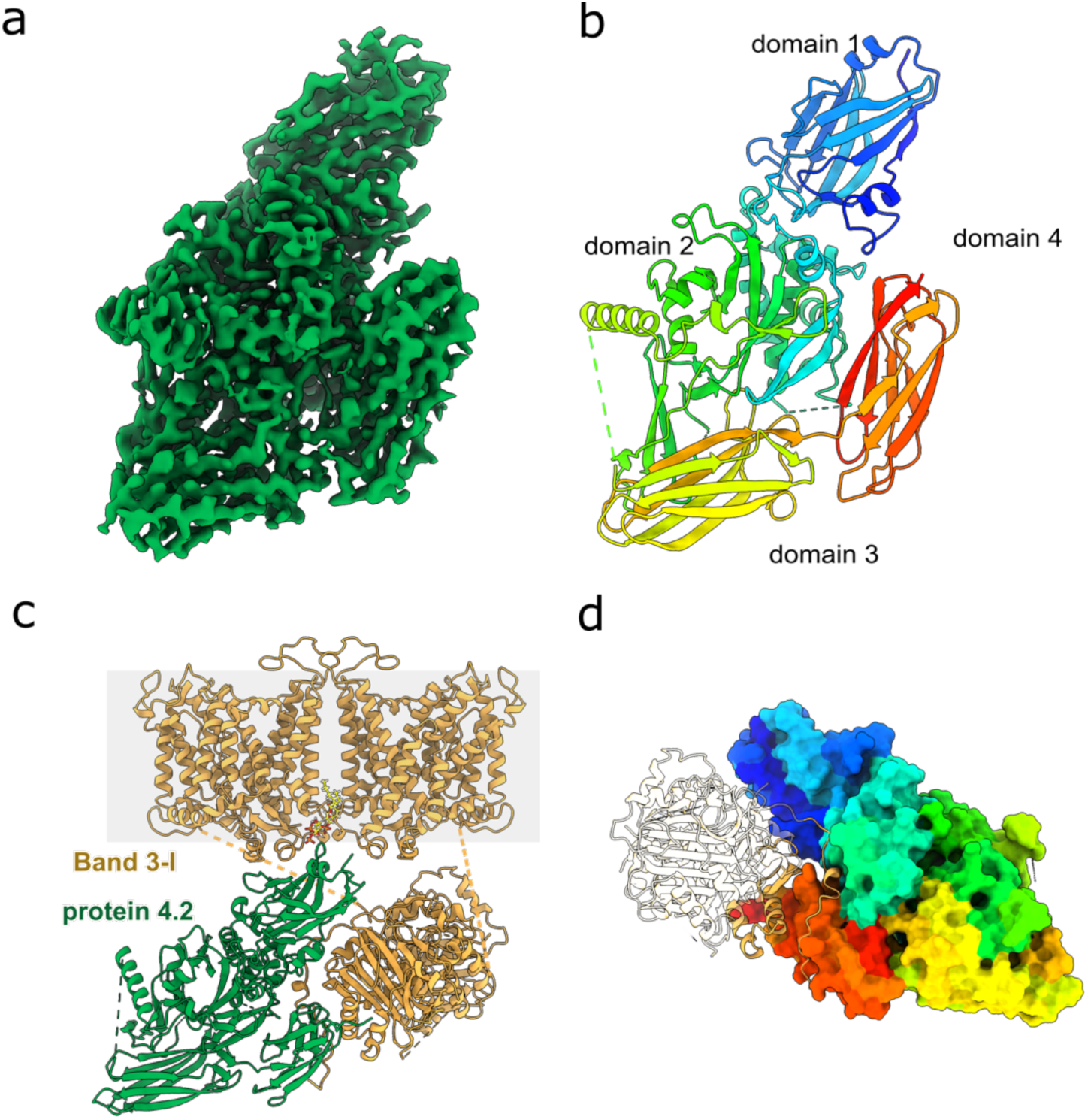
Protein 4.2 is an adaptor that mediates binding of Band 3-I. **a,** The 2.4 Å cryo-EM density map of protein 4.2 from local refinement of the consensus reconstruction. **b,** Structure of protein 4.2 colored by domain from the N- (blue) to the C- (red) terminus. The four domains are labeled. The proteins are depicted in cartoon representation. **c,** Model of protein 4.2 (green) and band 3-I (orange), PIP_2_ is colored in yellow. The proteins are showin in cartoon representation, and the membrane is represented in gray. **d,** The outer face of protein 4.2 forms the primary binding site for Band 3-I. Protein 4.2 is displayed as a molecular surface colored by domain from the N- (blue) to the C- (red) terminus, band 3-I cytosolic domain is displayed as ribbon, and the parts of Band 3-I that are not interacting with protein 4.2 are shown in transparency.

**Fig. 4:**
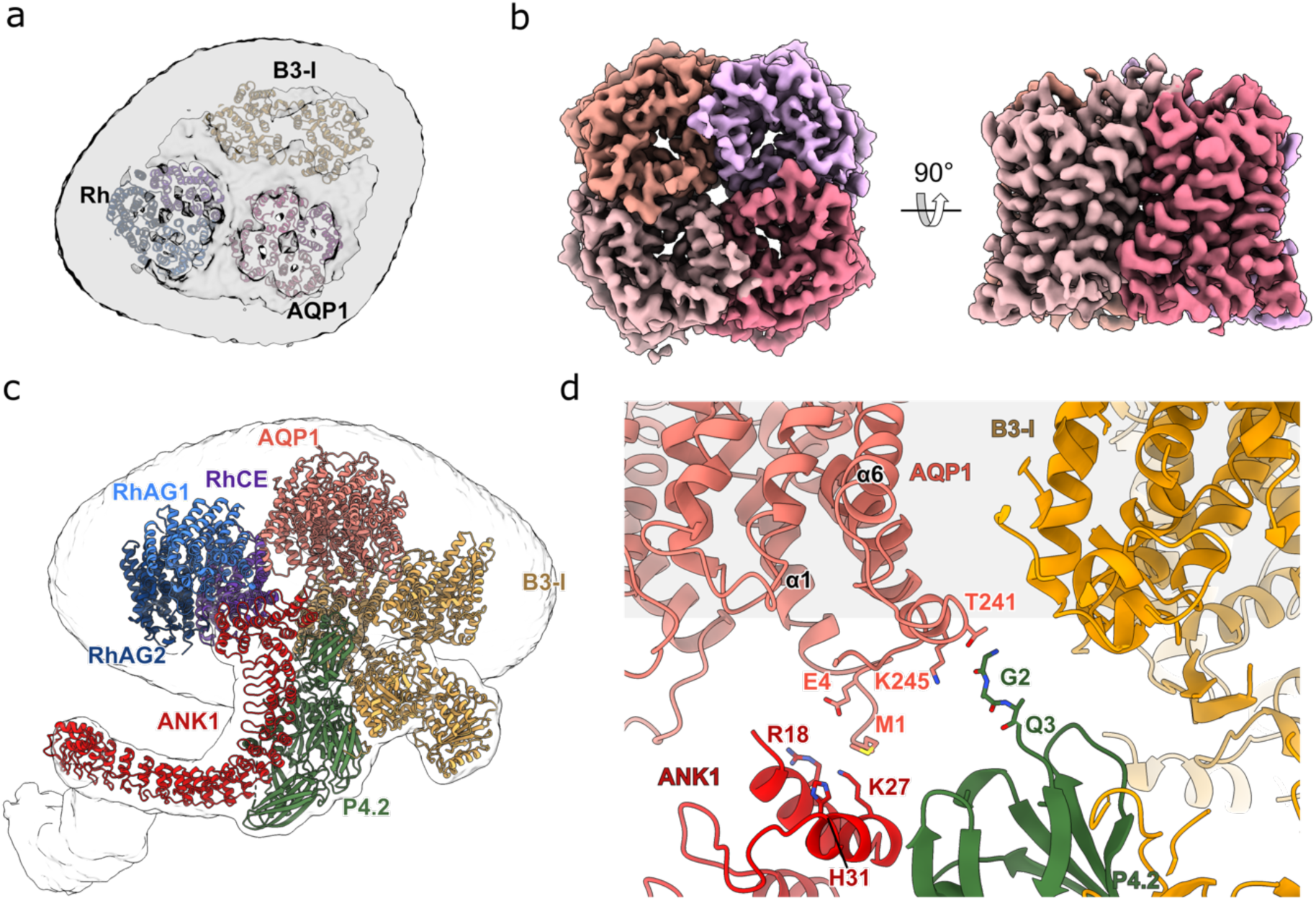
Aquaporin is a component of the ankyrin-1 complex. **a,** Aquaporin-1 is present in Classes 2 and 5 (Extended Data Fig. 5). The cryo-EM density map of Class 2 shows inside the membrane the density and atomic models corresponding to aquaporin (salmon), Rh (purple, blue and light blue) and Band 3-I (orange). **b,** The 2.8 Å cryo-EM density map of the aquaporin in Class 2. Densities corresponding to the four subunits are colored in magenta, deep pink, medium purple and hot pink. **c,** The model of Class 2 shows that AQP1 (salmon color) is located at the mutual interface of protein 4.2 (green), ankyrin (red), RhCE (purple) and Band 3-I (orange). **d,** Close up of the interaction region between AQP1, protein 4.2, ankyrin and Band 3-I. The membrane is represented in gray. The key residues that mediate the interactions in the interfaces are shown as sticks.

Refinement of all 850k ankyrin-1-containing particles against an *ab initio* model yielded a consensus map with an overall resolution of 2.6 Å (**Extended Data Fig. 3**). In the ankyrin-1 consensus reconstruction, well-ordered density is observed for the Rh heterotrimer, ankyrin, protein 4.2, and the cytoplasmic domains of a single band 3 dimer (denoted subsequently as Band 3-I). Masked refinement was used to improve the density quality for each component, giving a 2.4 Å reconstruction of RhAG/CE, a 2.6 Å reconstruction of ankyrin-1 (AR1-17), a 2.4 Å reconstruction of protein 4.2, and a 2.6 Å reconstruction of the cytoplasmic domains of band 3-I (**Extended Data Fig. 4**).

Sub-classification of the ankyrin-1 consensus refinement allowed us to identify six distinct classes with variable composition (**Extended Data Fig. 5a**). The core architecture described above is conserved in all six classes, but with elaborations. Class 1 had a larger micelle, accommodating two additional band 3 dimers (denoted Band 3-II and Band 3-III) that interact directly with ankyrin. 2D classes of Class 1 resemble complexes identified in native membrane vesicles using cryo-electron tomography (**Extended Data Fig. 6** and **Movie S1**). Classes 2 and 5 had an additional component present that proved to be an aquaporin 1 (AQP1) tetramer. Class 4 has an overall architecture similar to Class 1, but with a somewhat different orientation of the Band 3-I transmembrane region, and the presence of an additional membrane protein of unknown identity, denoted “X” herein. We have not assigned this component, but given the size of the density and its proximity to protein 4.2 and RhAG, we believe it most likely corresponds to CD47, the thrombospondin receptor, which is known to associate with both protein 4.2 and RhAG.

**Fig. 5:**
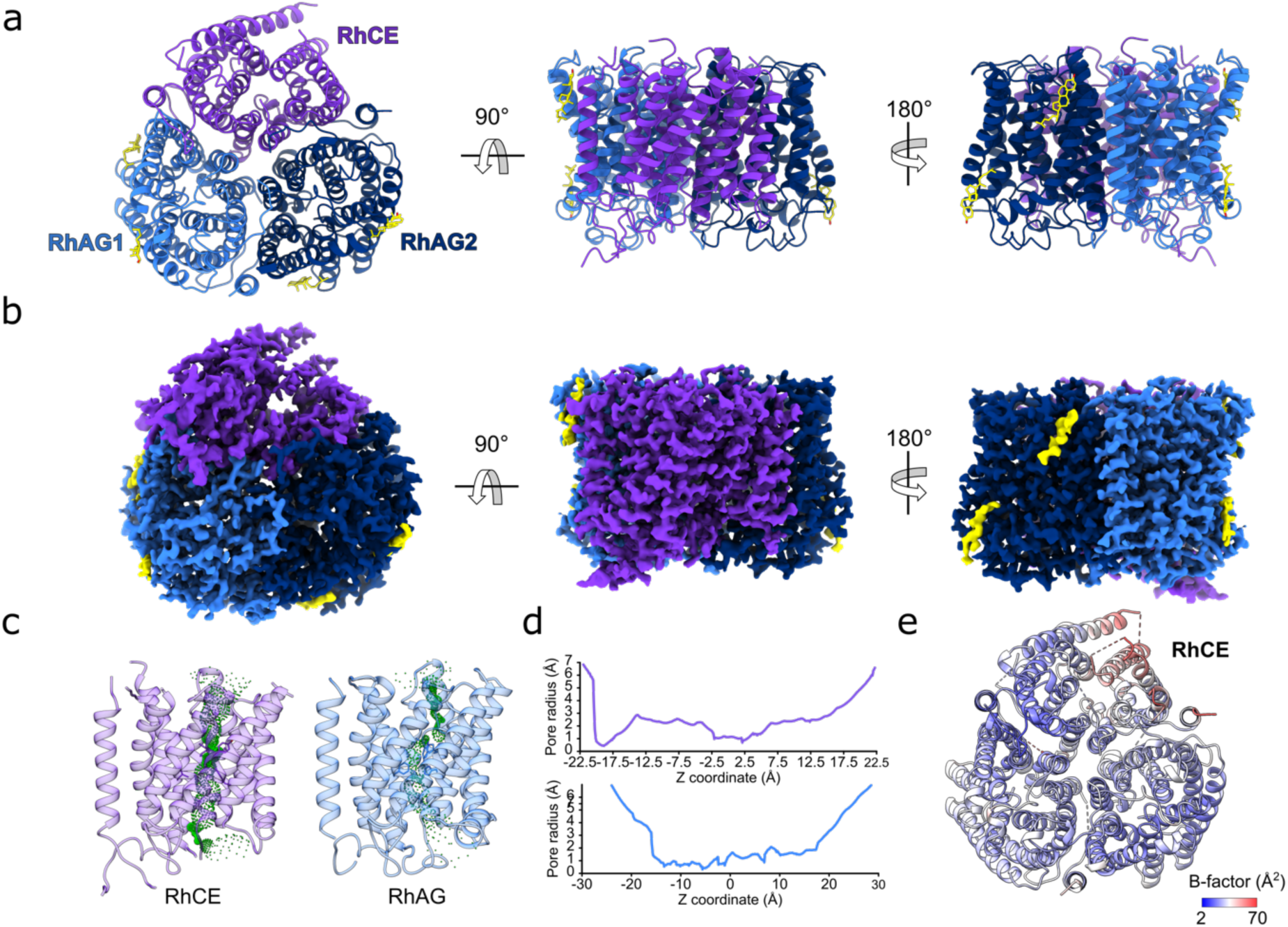
(RhAG)*_2_*(RhCE) trimer. **a,** The structure of the (RhAG)_2_(RhCE) trimer as viewed from the cytoplasm (right panel), and as viewed in the plane membrane (central and right panels). RhCE is colored in purple, RhAG1 in light blue and RhAG2 in dark blue. Two molecules of cholesterol (yellow) are bound to each RhAG. **b,** The 2.4 Å cryo-EM density map of the (RhAG)_2_(RhCE) complex. Densities corresponding to RhCE, RhAG1 and RhAG2 and cholesterol are colored as in panel A. **c,** Analysis of the pores of RhCE and RhAG. Figure of the pore in green generated by HOLE. **d,** 2D graph of pore radius for RhCE (top graph) and RhAG (bottom graph). **(E)** Refined atomic B-factors, averaged per residue. RhCE has higher B-factors than RhAG, likely indicating more flexibility/mobility.

**Fig. 6:**
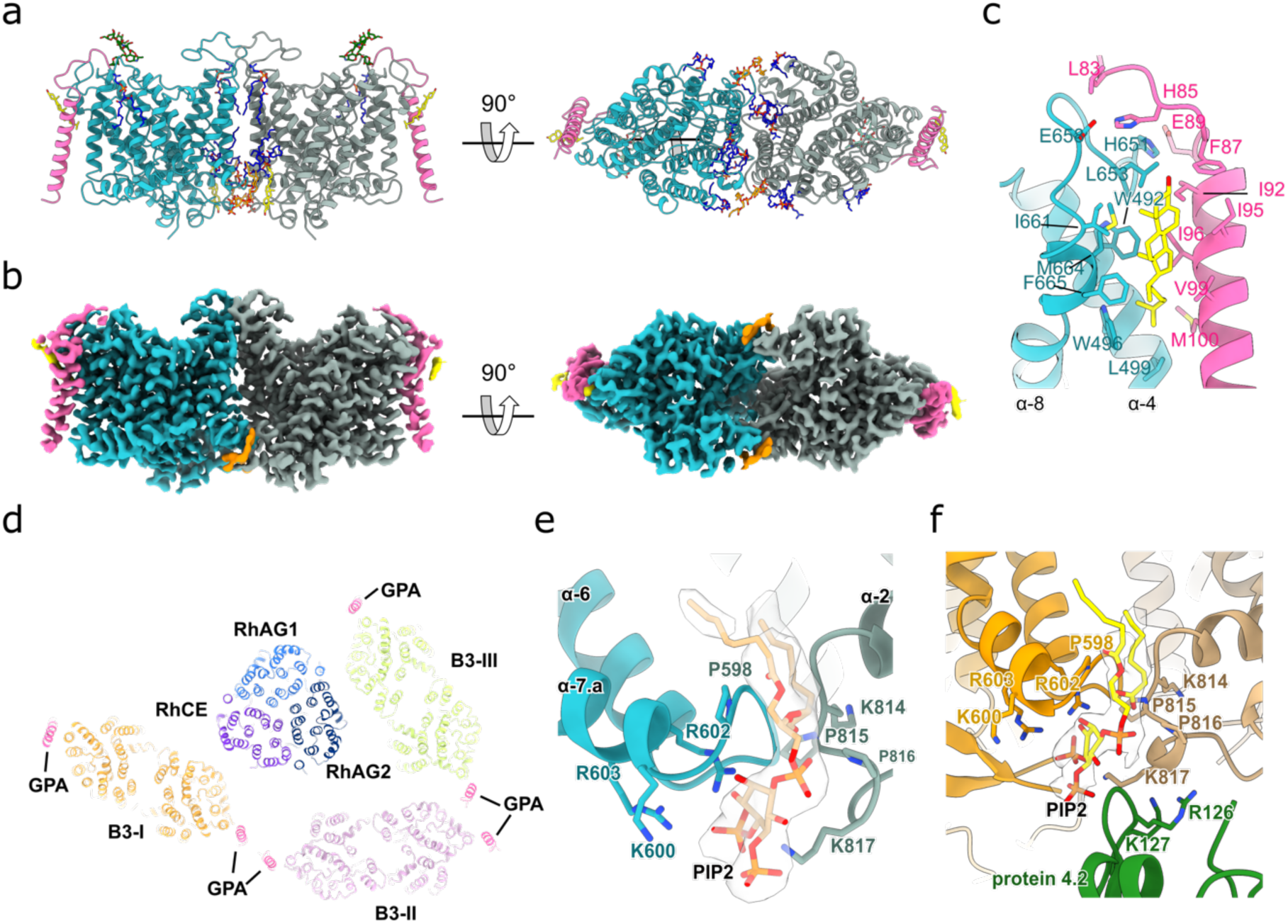
Band 3 in complex with glycophorin A. **a,** The structure of band 3 in complex with glycophorin A as viewed in the plane of the membrane, as viewed from the cytoplasm (central panel), and as viewed from inside the cell. Band 3 protomers are colored in aqua green and gray, glycophorin A in magenta, and molecules of cholesterol (yellow) PIP_2_ (orange) and POPC (blue) are shown. Glycans are colored in green. **b,** The 2.8 Å cryo-EM density map of the band 3- glycophorin A complex. Density corresponding to band 3, glycophorin, cholesterol (bound to glycophorin) and PIP_2_ are colored as in panel A. The three orientations of the map are the same as panel A. **c,** Close up of the interactions between band 3 (aqua green), glycophorin (magenta) and a cholesterol molecule (yellow). The key residues that mediate the interactions are shown as sticks. **d,** Cut through the TM of ankyrin-1 complex shows the position of glycophorin A bound to band 3 in the context of the complex. **e,** Close up of the band 3 transmembrane dimer interface. PIP_2_ (orange) sits in the middle of the site. The key residues that mediate the interactions in the interfaces are shown as sticks. **f,** PIP_2_ (orange) sits in the middle of the dimerization site of Band 3-I and close to the interaction site between Band 3-I and protein 4.2. The key residues that mediate the interactions in the interfaces and between Band 3-I TM and protein 4.2 are shown as sticks.

While the cytoplasmic domains of Band 3-I were well ordered in the consensus refinement, the transmembrane domains were very poorly resolved. In order to improve the density in this region, we applied the following strategy. Briefly, after local refinement of the transmembrane region of Band 3-I starting from the consensus refinement gave a poor quality 4.5 Å map, 3D-variability analysis was performed in cryoSPARC using a mask around the Band 3-I transmembrane region to identify modes of conformational and compositional heterogeneity. One mode appeared to correspond to an order-disorder transition of the transmembrane region. Clustering the particles using only this mode allowed identification of set of 108k particles with greatly improved density for the transmembrane region; local refinement of this particle set gave a 3.3 Å map with excellent density quality, and a well-resolved interface with protein 4.2 (**Extended Data Fig. 7**). A similar approach was applied to improve the density of Band 3-III starting from Class 1, resulting in a 3.8 Å local refinement.

By applying the 3D-VA-based clustering approach described above to Class 2, a subset of particles was identified with improved density for AQP1, resulting in a 3.7 Å C1 local refinement that facilitated unambiguous fitting of the AQP1 crystal structure, and identification of the regions of AQP1 that interact with ankyrin-1 and protein 4.2. Application of local C4 symmetry to the AQP1 tetramer yielded a 2.8Å map.

We have used the reconstructions described above to build atomic models for Class 1, which we believe represents an intact complex, and Class 2, in which the interaction with AQP1 is best defined. We assigned AQP1 in classes 2 and 5 due to the characteristic fold, which at 3.7 Å allows unambiguous fitting of the crystal structure of AQP1 to the density map. The remaining classes may represent genuine diversity in the composition of complexes present in the membrane, or partial dissociation of a single native complex (**Extended Data Fig. 5a**). It does not appear that the bound configuration of AQP1 in classes 2 and 5 would be incompatible with the arrangement of band 3 dimers in Class 1 (**Extended Data Fig. 5c**). We therefore assess it likely that *in vivo*, the three band 3 dimers identified here, as well as AQP1, are part of the same complex.

### Architecture of the human erythrocyte ankyrin-1 complex

The architecture of Class 1, the highest resolution intact complex (overall resolution 3.0 Å), is presented in Figure 1. The membrane-embedded core of the complex is formed by the Rh heterotrimer, consisting of a single RhCE protomer which forms the attachment site for ankyrin, and two RhAG protomers.

Ankyrin directly binds to the cytosolic N- and C-termini of RhCE, via the first five ankyrin repeats (AR1-5), which are oriented parallel to the membrane. The subsequent ankyrin repeats (AR6-24) emerge perpendicularly downwards from a T-interface with AR1-5, and gradually recurve to run almost parallel to the membrane plane.

Protein 4.2 is oriented vertically, such that the myristoylated N-terminus contacts the membrane, with one flat surface forming an extensive interface with AR1 and AR6-13 of ankyrin, while the distal face is available for recruitment of band 3.

Three band 3 dimers, each associated with two molecules of glycophorin A (GPA), form part of the complex. We have named them Band 3-I, Band 3-II and Band 3-III for the purpose of clarifying their distinct locations and environments. Band 3-I binds to the distal face of protein 4.2 via the band 3 cytoplasmic domain and N-terminal peptide. Band 3-II and Band 3-III, by contrast, both interact directly with ankyrin, but at distinct sites. Band 3-II interacts with the outer face of AR17- 19 via the cytoplasmic domain, while the cytoplasmic domain of Band 3-III binds to the outer face of AR21-24, and the Band 3-III N-terminal peptide runs back along the inner groove of ankyrin.

### Membrane association and target recruitment by ankyrin-1

We observe simultaneous engagement of the ankyrin-1 membrane binding domain with multiple targets (**Fig. 2**) that fall into three distinct categories: membrane embedded (RhCE & AQP1), extramembrane (cytosolic domains of band 3) and peripheral adaptors (protein 4.2). This is consistent with the previously described roles of all three vertebrate ankyrins in mediating spatial clustering of membrane proteins, for example facilitating clustering of NaV channels at the nodes of Ranvier^18^. The three categories of targets identified here interact with ankyrin-1 at distinct sites. The integral membrane targets, RhCE and AQP1, both interact with the peptide binding groove of AR1-5; the adaptor (protein 4.2) interacts with the exterior surface of AR1-13; and the two extramembrane targets (Band 3-II & Band 3-III) interact predominantly with the exterior surface of AR17-24, supplemented in the case of Band 3-III by interactions of the N-terminus with the peptide binding groove of AR6-10.

The Rh heterotrimer forms the primary membrane attachment site for ankyrin-1, mediated via an interaction between the N- and C-termini of RhCE and the first five repeats of ankyrin-1 (**Fig. 2e**). The structure of ankyrin-1 in the complex (**Extended Data Fig. 8a**) is markedly distinct from that expected based on structures of the AR1-24 and AR1-19 fragments of ankyrin-2 bound to autoinhibitory motifs^19^. The first repeats are dramatically rearranged compared to their position in the ankyrin-2 structures (**Extended Data Fig. 8b**). The structures of ankyrin-2 were determined in the absence of a membrane embedded target protein, and the presence of an ankyrin-1 autoinhibitory motif, suggesting the possibility that ankyrin-1 may adopt a similar conformation in solution, converting to the T-shaped arrangement only upon engagement of an integral membrane target. Flexibility at this interface is not entirely unexpected; comparison of the structure of AR1-9 of ankyrin-2 bound to an NaV channel peptide^20^ with the aforementioned AR1- 19 and AR1-24 structures bound to the autoinhibitory peptide shows substantial flexibility at this interface (**Extended Data Fig. 8c**), albeit not quite to the extent seen here. A sequence alignment of the 24 repeats of ankyrin-1 (**Extended Data Fig. 8d**) shows that, while AR5 and AR6 are highly conserved when compared to the equivalent regions of ankyrin-2 and ankyrin-3 (**Extended Data Fig. 8f**), they are the most divergent of the ankyrin repeats when compared with the other repeats in the ankyrin-1 MBD. Notably, AR5 is shorter than the other repeats by 4 residues, and AR6 lacks both the “S/TP” motif at positions 4-5, and the consensus “GH” motif at positions 13-14 is replaced by two aspartates. The rearrangement of the AR1-5 module orients the canonical protein binding groove to directly face the membrane, in order to bind the membrane-embedded targets RhCE (in all classes) and AQP1 (Classes 2 & 6) (**Extended Data Fig. 5a**).

The flat surface of protein 4.2 is oriented vertically with respect to the membrane, and forms multiple sites of attachment to ankyrin-1, all of which are outside the canonical peptide binding groove (**Fig. 2f**). Protein 4.2 acts as an adaptor molecule and a stabilizer for ankyrin, with the other face of protein 4.2 forming the attachment site for Band 3-I (**Fig. 2b**). Interaction of domain 1 of protein 4.2 with the exposed “edge” of AR1 appears to further stabilize the T-shaped conformation of the ankyrin-1 membrane binding domain (**Fig. 2f**).

In Class 1, Band 3 -II and Band 3-III directly bind to ankyrin, interacting with repeats 17-19 & 21- 24 respectively (**Fig. 2c-d**). The N-terminus of Band 3-III runs back along the inner ankyrin groove, with residues 2-24 forming an ordered interaction with AR6-10 and the AR5-6 linker (**Extended Data Fig. 9**). Notably, this peptide is oriented in the reverse direction to that previously observed for ankyrin autoinhibition motifs (**Extended Data Fig. 9d**), but in the same direction observed for the binding of the NaV 1.2 N-terminal peptide to repeats AR1-9 of ankyrin-2^20^ (**Extended Data Fig. 10e**). The orientation of the Band 3-I cytoplasmic domains is inverted with respect to the membrane when compared with Band 3-II & Band 3-III, which are both directly bound by ankyrin (**Extended Data Fig. 10d**). Notably, interacting surfaces of band 3 cytoplasmic domains are distinct and non-overlapping (**Extended Data Fig. 10c**).

### Structure of protein 4.2 and interactions with band 3

The structure of the erythrocyte ankyrin-1 complex also represents the first experimentally determined structure of protein 4.2, solved here at a resolution of 2.4 Å (**Fig. 3a-b**). As previously suggested^21^, the fold of protein 4.2, comprising four compact domains assembled in a flat, P-shaped arrangement, is very similar to that of the closed conformation of tissue transglutaminase (TG2). The central papain-like domain (domain 2) is surrounded by three Ig-like domains (domains 1, 3 & 4). The N-terminal glycine of protein 4.2 is known to be myristoylated^22^, and indeed in the present structure we see that the N-terminal glycine of protein 4.2 directly contacts the membrane. The vertical orientation of protein 4.2 exposes two flat sides, which serve to mediate protein-protein interactions. The inner face interacts with ankyrin-1, as described in the previous section, while the outer face forms the primary binding site for Band 3-I (**Fig. 3c-d**). The Band 3-I binding mode shares some broad similarities with the binding mode of Band 3-II to ankyrin (**Extended Data Fig. 10d-e**), insofar as it involves recognition of a 3D epitope via an extensive interface, and augmentation by interaction of the free N-terminus of band 3 with a peptide recognition groove. In the case of protein 4.2, the interaction with Band 3-I is mediated by interactions with D4 & D1 of protein 4.2 (**Fig. 2b**), and the N-terminus of Band 3-I is nestled in a groove formed by D1, D2 & D4. In Class 2, D1 of protein 4.2 also appears to be forming interactions with AQP1, although the structural details of this are unclear, due to limited local resolution in the AQP1 tetramer. D1 is also forming an interface with loops near the transmembrane dimerization interface of Band 3-I, in a region that also forms cytoplasmic-TM contacts in Band 3-II (**Extended Data Fig. 10e**).

### Aquaporin is a component of the ankyrin-1 complex

AQP1 was not previously known to participate in the erythrocyte ankyrin complex, although it has been previously shown to co- localize within 80 Å of a population of band 3 dimers using FRET spectroscopy^23^. A final resolution of 3.7 Å allows unambiguous fitting of the crystal structure of AQP1 to the density map (Fig. 4B). AQP1 is located at the mutual interface of Band 3-I, protein 4.2, and AR1-2 (**Fig. 4a- c**). The interaction appears to be largely mediated by interactions of the N-terminus of AQP1 with AR1-2, and interactions of the C-terminus of AQP1 with domain 1 of protein 4.2 (**Fig. 4d**). In addition, strong, unmodeled density is present in the cleft between AQP1 and RhCE, suggesting the possibility of a lipid-mediated interaction with the Rh heterotrimer (**Extended Data Fig. 11g**). Although the identification of aquaporin in the ankyrin complex was initially surprising, it is consistent with previous studies showing that hereditary spherocytosis patients have a reduction in AQP1 expression and plasma membrane localization^24^. Furthermore, studies of erythrocytes in a band 3 knockout mouse model show that AQP1 is reduced in band 3 (-/-) erythrocytes, and has a distinctly altered glycosylation pattern, suggesting perturbed trafficking in the absence of band 3^25^.

### Structure and composition of the Rh channel

The heterotrimeric Rh ammonia channel plays an important role in ammonium homeostasis in the blood^26^, and in maintaining the stability of the red blood cell membrane, possibly via direct interaction with ankyrin-1^17^. Humans express five Rh family proteins – RhAG, RhCE and RhD in erythrocytes, and RhBG and RhCG in the kidney epithelium. The structure of the RhCG homotrimer ^27^ showed that the eukaryotic Rh family shares a great deal of similarity to the bacterial AmtB family of ammonium transporters, including a conserved twin histidine motif in the central pore. Based on the absence of this twin histidine motif in the sequence of RhCE and RhD, it has been suggested that RhAG mediates ammonia transport, while RhCE/D play some as yet undefined role^28^. The stoichiometry of the channel has also remained uncertain.

In the structure of the erythrocyte ankyrin-1 complex presented here, we have resolved the structure of the Rh heterotrimer at 2.4 Å, (**Fig. 5a-b**, and **Extended Data Fig. 4**) revealing that two molecules of RhAG and one molecule of RhCE form the heterotrimeric channel (**Extended Data Fig. 11a**), with the N- and C- termini of RhCE forming the primary interface with ankyrin- 1 (**Fig. 2e**) Several well-ordered cholesterol molecules are bound to the transmembrane regions of both RhAG and RhCE on both the inner and outer leaflets of the membrane (**Fig. 5a-b**). The pore of RhAG shows the expected twin-histidine motif in the center of the pore (**Extended Data Fig. 11b**), with a hydrogen bonded network of water molecules similar to what has been observed for AmtB and RhCG. The pore of RhCE is blocked at a short hydrophobic constriction; however, the presence of matched cavities on both the cytosolic and extracellular sides extending almost completely across the membrane (**Fig. 5c-d**) raises the possibility that this subunit may have some yet to be elucidated role in membrane transport. The density for RhCE is also weaker than for RhAG, despite the fact that RhCE is the subunit that directly engages with ankyrin. This weaker density (and correlated higher atomic B-factors) may indicate flexibility, possibly consistent with an unresolved conformational change corresponding to pore opening (**Fig. 5e**).

The source of our sample was derived from RhD positive donors, and therefore contains both RhCE & RhD. Given the close homology of RhD and RhCE, which vary at only 35 positions, this raises the question of whether RhCE, RhD, or a mix of the two is present in the structurally resolved complex. Analysis of the model-map fit at the sites of variation (**Extended Data Fig. 11d**) strongly suggests that RhCE is predominantly incorporated, consistent with previous findings suggesting preferential association of RhCE with the band 3 complex^29^, although we cannot exclude the possibility that a small fraction of RhD is present; extensive attempts at focused classification in RELION 3.1^30^ did not allow isolation of a class corresponding to RhD-containing trimers. This is consistent with previous genetic studies showing that RhD and RhCE are not interchangeable in the context of the ankyrin-1 complex, and that absence of RhCE has significant effects on the composition of the complex^31^.

The interaction of RhCE with ankyrin is mediated in large part by the binding of the N- and C- termini of RhCE to the peptide binding groove of AR1-5 of ankyrin-1 (**Fig. 2e**). RhAG also helps stabilize this interaction, as the RhCE termini pack on top of the adjacent RhAG protomers. Key residues in the RhCE and RhD termini that mediate the binding of ankyrin are not conserved in RhAG, suggesting that RhCE/D most likely form the exclusive ankyrin binding interface in erythrocytes

### Structure of the band 3 dimer complexed with glycophorin A

In the structure of the free band 3 dimer (**Fig. 6a-b**), a glycophorin monomer is bound at each end of the complex, with the transmembrane helix interacting primarily with TM4 of band 3, and the extracellular N-terminal peptide interacting with the TM7-8 loop (**Fig. 6c**). A tightly bound cholesterol molecule is located near the extracellular side, wedged between the transmembrane helix of glycophorin A and TM8 of band 3 (**Fig. 6c**). A prominent lipid headgroup density is located at the intracellular dimer interface of band 3, where it binds to a patch of positively charged residues. Based on the distinctive shape of the density, we have assigned this density as PI(4,5)P_2_ (**Fig. 6e**). This density is also observed in classes of the intact ankyrin complex, where it is located very close to the interface of Band 3-I with protein 4.2 (**Fig. 6d,f**). Only monomeric GPA is observed binding to band 3, despite the known propensity of GPA to dimerize. This is consistent with studies showing that mutations which inhibit GPA dimerization do not inhibit band 3 dependent effects of GPA^32^, but still leaves a puzzle: why does the GPA dimer not interact with band 3? Structural alignment of the crystal structure of the GPA dimer ^33^ with band 3-bound GPA shows that the second molecule of GPA would have to be embedded entirely within the membrane, in an almost horizontal orientation, which is highly energetically unfavorable. We speculate that in the membrane, GPA exists in a monomer-dimer equilibrium, and band 3 selectively binds monomeric GPA. In the context of the Class 1 model of the ankyrin-1 complex, the GPA molecules bound to Band 3-II and Band 3-III are in close proximity, though we cannot be certain they are interacting. The possibility of GPA mediating end-to-end attachment of two band 3 dimers would be consistent with both previous molecular dynamics simulations of band 3 in the presence of modeled GPA^34^, as well as the known propensity of detergent-solubilized band 3 purified from erythrocytes to form “tetramers” in solution^35^.

## DISCUSSION

Ankyrins form part of an intricate sub-membrane lattice that provides mechanical stability, shape, and local functional organization to cellular membranes in many cell types including erythrocytes and neurons. In particular, in erythrocytes, ankyrin-1 forms the nucleus of a macrocomplex which acts to tether spectrin to the membrane ^3^ and organizes several highly abundant membrane proteins involved in gas exchange, pH regulation, and control of cellular volume^25^. Mutations in elements of this complex lead to defects in erythrocyte size, shape and stability^15^. However, the architecture and composition of the complex, as well as the mechanism of ankyrin-1 recruitment to the membrane, have so far remained unresolved. Whether the so-called “Rh antigen complex” and “band 3 macrocomplex” are distinct, separate complexes which interact, or in fact one single supercomplex, has also been a matter of debate. Here, we used single-particle cryo-EM to determine structures of an erythrocyte ankyrin complex that includes both the RhAG/CE heterotrimer and band 3. We provide a mechanistic hypothesis for ankyrin recruitment and autoinhibition, showing how ankyrin-1 can recruit three band 3 dimers at distinct sites, clarifying the role of protein 4.2, and identifying aquaporin-1 as an unanticipated component of the ankyrin- 1 complex.

### Membrane-bound ankyrin-1 adopts a T-shaped conformation

The discontinuous, T-shaped conformation of ankyrin-1 we observe in the membrane bound state is strongly suggestive of a rearrangement upon membrane binding, which we propose is linked to the mechanistic basis of both ankyrin membrane association and autoinhibition. Thus far, the only available structure of a full-length ankyrin membrane binding domain is that of ankyrin-2 fused to the autoinhibitory C-terminal tail of ankyrin-1^19^, which binds in a continuous groove that traverses the entire length of the molecule. Constructs lacking the autoinhibitory peptide failed to crystallize, likely because they were too conformationally heterogeneous. In contrast, in our structure (**Fig. 1a**), the ankyrin- 1 repeats AR1-5 are oriented parallel to the membrane, and orthogonal to the downstream ankyrin repeats. While there is no comparable structure of ankyrin-1 available, the membrane binding domain and the AR5-6 region in particular are highly conserved in all three ankyrin proteins, suggesting that this is unlikely to be an isoform-specific difference. In the ankyrin-2 AR1-24 structure, the autoinhibitory peptide occupies a groove that crosses the AR5-6 boundary, stabilizing them in the “connected” state. By contrast, in our structure, the N-terminal peptide of Band 3-III occupies this same groove, but ends at the AR5-6 interface, with the very N-terminal 11 residues, including the Syk phosphorylation site Y8, appearing to stabilize it in the T-shaped configuration (fig. S9, D and E). Deletion of these 11 residues causes a severe hemolytic anemia, without compromising insertion into the membrane^36^. Phosphorylation of Y8 also causes alterations in erythrocyte size and shape^37^, consistent with a role for this segment in stabilizing the membrane associated configuration of ankyrin. We hypothesize that the band 3 N-terminal peptide displaces the autoinhibitory motif from the inner groove of ankyrin, freeing AR1-5 to adopt the membrane associated configuration, and leaving the AR1-5 groove unoccupied, facilitating recognition of RhCE and AQP1. Binding to protein 4.2 and RhCE further stabilizes the T-shaped configuration of ankyrin, resulting in the stable core complex we observed here. In this model, the autoinhibited state is not competent to bind to the membrane not only because the inner groove is occupied, as has been suggested previously, but also because AR1-5 are in a conformation that cannot orient the groove to bind membrane embedded targets.

### Band 3 clustering mediated by ankyrin-1 and protein 4.2

The ankyrin-1 complex structure reveals four distinct sites of target recruitment (**Fig. 2a**). We term these sites the juxtamembrane (AR1-5 binding of RhCE and AQP1), the adaptor-mediated (protein 4.2 recruitment of Band3-I), the central (AR16-18 binding of Band3-II) and the distal site (Band-3-III binding, including interactions of N-terminal peptide). The observation that band 3 associates directly with ankyrin at the distal and central sites is consistent with studies of AR13-24 binding to the cytoplasmic domain of band 3, which identified a high affinity and a low affinity site^38^. Band 3 has variously been characterized as binding to ankyrin as a bona fide tetramer, or a “pseudotetramer”, in which ankyrin links two band 3 dimers. Our structure is consistent with the latter model. The central site, is broadly in agreement with the EPR-based structural model of Kim et al.^39^, as well as studies showing the importance of loops 175-185 and 63-73 of the band 3 cytoplasmic domain for interaction with ankyrin. In contrast, the distal site uses an entirely different interface on both ankyrin and the band 3 cytoplasmic domain, and is further stabilized by interactions of the N-terminal peptide of band 3 with the peptide binding groove of ankyrin. The interaction of the N-terminus of band 3 with the ankyrin-1 peptide binding groove occurs at the same site, but with the opposite directionality, as has previously been characterized for the interaction of ankyrin autoinhibitory peptides^19^. In addition to the ankyrin-mediated association of Band 3-II and Band 3-III via their cytoplasmic domains, the proximity of the glycophorin A molecules bound to Band 3-II and Band 3-III suggests the possibility of glycophorin A mediated end-to-end contacts between the transmembrane domains, as suggested by molecular dynamics simulations^34^.

### Protein 4.2 acts as an adaptor, mediating the recruitment of Band 3-I and AQP1

The structure of the ankyrin-1 complex also establishes a role for protein 4.2, a highly abundant component of the red blood cell. Protein 4.2 has been previously shown to interact with both band 3 and ankyrin^40^, but the implication of these interactions was not clear. We observe that protein 4.2 plays a role as an adaptor, stabilizing the interaction of ankyrin with the membrane and facilitating recruitment of an additional dimer of the band 3 anion exchanger (Band 3-I) via interactions with both the cytoplasmic and TM regions. The latter of which appears to be at least partially mediated by a PIP2 lipid bound at the band 3 transmembrane dimer interface. Furthermore, protein 4.2 appears to play an additional role in facilitating the recruitment of AQP1, the C-terminus of which directly interacts with domain 1 of protein 4.2.

### Aquaporin-1 is a component of the ankyrin-1 complex

The identification of aquaporin- 1 as a component of the ankyrin-1 complex (Fig. 4) was an unexpected finding, although consistent with FRET measurements indicating that AQP1 is located within 8nm of band 3 dimers. The observation that AQP1 expression is decreased, and glycosylation pattern altered in both murine and human erythrocytes lacking band 3^25^ is also compatible with a role for AQP1 in the ankyrin- 1 complex. It has been proposed that the cluster of membrane proteins that ankyrin-1 organizes in the red blood cell membrane forms a “CO_2_ metabolon”^41^, linking facilitated diffusion of CO_2_ across the membrane, anion exchange of Cl^-^/HCO_3-_ via band 3, and interconversion of dissolved CO_2_ and carbonic acid in the cytosol via carbonic anhydrase, which binds to the C-terminal tail of band 3^42^ and catalyzes conversion of CO_2_ and H_2_O to HCO_3-_ and H^+^. A role for AQP1 in the CO_2_- metabolon would be consistent with the known capacity of AQP1 to conduct both CO_2_ and H_2_O ^43^, both of which are substrates for carbonic anhydrase. RhAG may also participate indirectly in this process, by allowing facilitated diffusion of NH_3_ and consequent modulation of [H^+^], via the equilibrium of NH_3_ + H^+^ and NH_4_^+^.

### RhCE as an ankyrin-1 recruitment site and a putative orphan solute channel

The functional role of the homologous proteins RhCE and RhD in the structure and function of the red blood cell membrane has been much debated, as has the stoichiometry of the heterotrimeric Rh complex. The 2.4 Å resolution structure of the RhAG2RhCE heterotrimer (**Fig. 5a-b**) in the context of the ankyrin-1 complex resolves the stoichiometry question, and suggests that the primary role of RhCE might be as a recruitment site for ankyrin-1, via binding of the N and C-termini to AR1-5 of ankyrin-1. Other putative roles for RhCE remain speculative. It has been proposed, based on homology modeling^44^ and the absence in RhCE/D of the characteristic twin histidine motif which is associated with ammonium and possibly carbon dioxide transport in RhAG, RhBG and RhCG, that RhCE and RhD lack a membrane transport function completely. Our work, while not conclusively demonstrating that RhCE retains a transport function, shows that RhCE has a pore which extends almost completely across the membrane, impeded by only by a small cluster of hydrophobic residues. This pore has entries on both the extracellular and intracellular sides of the membrane, and based on inspection of the density map and refined atomic B-factors of the model, appears more dynamic than those of the two RhAG subunits. Taken together, we suggest that the possibility of an auxiliary transport function for RhCE in the red blood cell membrane may be worth investigating.

### Concluding remarks

In summary, we have reported the structure of an erythrocyte ankyrin-1 complex, purified from human erythrocytes. We have structurally characterized two direct binding sites for the band 3 anion exchanger on ankyrin-1, identified aquaporin-1 as a component of the complex, determined the stoichiometry of the heterotrimeric Rhesus ammonia channel, and identified RhCE as the ankyrin-1 binding subunit of the Rhesus channel. Taken together these findings offer insights into the molecular architecture of the red blood cell membrane, and show how ankyrins can facilitate the simultaneous recruitment of multiple divergent classes of membrane proteins, to enable functional organization of membrane transport processes.

## Methods

### Preparation of erythrocyte ghost membranes

Ghost membranes were prepared as described in^45^. Briefly, red blood cells from healthy blood donors were provided by the Transfusion Centre of the Hospital of Padua (Italy). Samples were obtained from voluntary and informed blood donation for transfusions. Erythrocytes from fresh human blood samples were washed twice in 5 volumes of 130 mM KCl, 10 mM Tris-HCl, pH 7.4. The cells were hemolyzed in 5 volumes of 1 mM EDTA, 10 mM Tris-HCl, pH 7.4, and centrifuged at 18,000 x g for 10 min. The ghost membranes were then washed five times in the hemolysis buffer, and four additional times in 10 mM HEPES, pH 7.4. The hemoglobin-free ghost membranes were finally resuspended in 130 mM NaCl, 20 mM HEPES, pH 7.4, 0.5 mM MgCl_2_, 0.05 mM CaCl2, 2 mM DTT, and stored at -80°C.

### Purification of the erythrocyte ankyrin-1 complex

The general purification workflow is presented in **Extended Data Fig 1**. Ghost membranes were solubilized at a protein concentration of 4 mg/ml in 130 mM KCl, 10 mM HEPES, pH 7.4, protease inhibitor tablet (cOmplete™, EDTA-free Protease Inhibitor Cocktail, Millipore Sigma), 1% (w/v) digitonin (Carbosynth), 1 mM ATP, 1 mM MgCl_2_, 1 mM PMSF, at 4°C for 1 hour. Unsolubilized material was removed by centrifugation at 26.000 x g for 30 min. The supernatant was applied to a PD10 column (to reduce the detergent concentration) equilibrated with 0.05% (w/v) digitonin, 130 mM KCI, 20 mM HEPES, pH 7.4, 1 mM ATP, 1 mM MgCl_2_, 1 mM PMSF. The sample was than applied on a glycerol step gradient (30-12% glycerol) and centrifuged for 15 h at 25,000 rpm (SW 32 rotor, Beckman). The separation and distribution of proteins was confirmed by SDS-PAGE gel (4–20% Mini-PROTEAN® TGX™ Precast Protein Gels, Biorad). Fractions containing high molecular weight species were pooled together and concentrated to <500 µL using a 100-kDa cutoff concentrator. The sample was loaded on a Superose 6 10/300 Increase column (Cytiva) equilibrated in 0.05% (w/v) digitonin, 130 mM KCI, 20 mM HEPES, pH 7.4, 1 mM ATP, 1 mM MgCl_2_, 1 mM PMSF. Fractions of a peak around 11 mL elution volume were analyzed using a mass photometer (Refeyn OneMP), pooled and concentrated to 8 mg/mL.

### Vesicle preparation

Ghost membranes (∼100 mg) were pelleted (6,000 x g, 10 minutes), washed 5 times in 130 mM KCl, 10 mM HEPES, pH 7.4, and then homogenized with the same buffer. The sample was sonicated using microprobe (amplitude = 30, pulse on = 5 sec, rest time = 10 sec, total pulse on = 50 sec). Large fragments are removed by centrifugation (6,000 x g, 10 minutes) and the supernatant was extruded (Avanti Polar Lipid) 10 times using a 400 nm filter. After extrusion, vesicles were collected by ultra-centrifugation at 35,000 rpm for 30 minutes (S120-AT3, Sorvall). The pellet, which contains vesicles, was resuspended at a final concentration of 5 mg/mL. All steps were performed at room temperature to facilitate vesicle formation.

### Mass photometry (MP, iSCAMS)

Mass photometry (MP) experiments were performed using a Refeyn OneMP (Refeyn Ltd.). Data acquisition was accomplished using AcquireMP (Refeyn Ltd. 172 v2.3). Samples were evaluated with microscope coverslips. The coverslips (Cover glass thickness 1 ½ 24 x 50 mm, Corning) were washed with ddH_2_O and isopropanol. Silicone templates were placed on top of the coverslip to form reaction chambers immediately prior to measurement. The instrument was calibrated using NativeMark Protein Standards (NativeMark™ Unstained Protein Standard, Thermo Fisher). 10 μL of fresh room temperature buffer was pipetted into a well, the focal position was identified and locked. For each acquisition 1 μL of the protein (at a final concentration after dilution of 200 nM) was added to the well and thoroughly mixed. MP signals were recorded for 60 s to allow detection of at least 2 × 103 individual binding events. Data analysis was performed using the DiscoverMP software.

### Cryo-EM & Cryo-ET grid preparation and data collection

3 μL of purified ankyrin complex at 8mg/mL, with a surface-active additive (0.01% (w/v) glycyrrhizic acid) was added to a glow discharged (PELCO easiGlow) holey gold grid (Quantifoil UltrAuFoil grids (R0.6/1, Au 300 mesh gold)) and blotted for 4 to 6 seconds at 4°C and 100% humidity using the Vitrobot system (ThermoFisher), before plunging immediately into liquid ethane for vitrification. Images were collected on a Titan Krios electron microscope (FEI) equipped with a K3 direct electron detector (Gatan) operating at 0.83 Å per pixel in counting mode using Leginon automated data collection software. Data collection was performed using a dose of 58 e-/Å2 across 50 frames (50 ms per frame) at a dose rate of 16 e–/pix/s, using a set defocus range of −0.5 μm to −1.5 μm. A 100 μm objective aperture was used. A total of 14464 micrographs were collected.

Vesicles for cryo-ET were prepared similarly using 1.2/1.3 μm holey gold grids (Quantifoil UltrAuFoil) and blotted for 10 s using a Vitrobot Mark IV (ThermoFisher), before plunging immediately into liquid ethane for vitrification. About 100 tilt series were collected on a Titan Krios electron microscope (FEI) equipped with a K3 direct electron detector (Gatan) and post-column Cs-corrector operating at 2.077 Å per pixel in counting mode using Leginon automated data collection software^46^. Tilt series were collected bi-directionally from [0:+51] degrees then [0:-51] degrees in 3 degree increments, with a dose of 3.13 e-/Å2 across 8 frames (57 ms per frame) total dose of 112 e-/Å2 across 36 total images per tilt series, using a set defocus range of −3.5 μm to −4.5 μm.

### Cryo-EM data processing

The final cryo-EM data processing workflow is summarized in Figure S3. Orientation distributions, FSC plots, and validation statistics are presented in **Extended Data Fig. 2** and **Table 1**. Maps, masks and raw movies have been deposited at EMDB (IDs: xxx) and EMPIAR (IDs: yyy). Subsequent steps were performed in cryoSPARC v3.2 unless otherwise indicated^47^.

**Table 1.** Cryo-EM data collection, refinement, and validation statistics.

|  | Ankyrin-1 complex<br>Class 1 | Ankyrin-1 complex<br>Class 2 | Ankyrin-1 complex<br>Class 3 | Ankyrin-1<br>complex<br>Class 4 |
| --- | --- | --- | --- | --- |
|  | (EMDB-TBA)<br>(PDB-TBA) | (EMDB-TBA)<br>(PDB-TBA) | (EMDB-TBA)<br>(PDB-TBA) | (EMDB-TBA)<br>(PDB-TBA) |
| **Data collection and processing** |  |  |  |  |
| Magnification | 105,000x | 105,000x | 105,000x | 105,000x |
| Voltage (kV) | 300 | 300 | 300 | 300 |
| Electron exposure (e-/Å<sup>2</sup>) | 58 | 58 | 58 | 58 |
| Defocus range (µm) | -0.5 to -1.5 | -0.5 to -1.5 | -0.5 to -1.5 | -0.5 to -1.5 |
| Pixel size (Å) | 0.83 | 0.83 | 0.83 | 0.83 |
| Symmetry imposed | C1 | C1 | C1 | C1 |
| Initial micrographs (no.) | 15,756 | 15,756 | 15,756 | 15,756 |
| Final micrographs (no.) | 14,464 | 14,464 | 14,464 | 14,464 |
| Initial particle images (no.) | 1,781,329 | 1,781,329 | 1,781,329 | 1,781,329 |
| Final particle images (no.) | 129,621 | 175,081 | 123,528 | 81,324 |
| Map resolution (Å) | 3.1 | 2.9 | 3.1 | 4.1 |
| FSC threshold |  |  |  |  |
| Map sharpening B factor (Å<sup>2</sup>) | 56.5 | 59.1 | 57.2 | 60.0 |
| Model composition |  |  |  |  |
| Nonhydrogen atoms | 126,050 |  |  |  |
| Protein residues | 7,760 |  |  |  |
| Ligands | 67 |  |  |  |
| B factors (Å<sup>2</sup>) |  |  |  |  |
| Protein |  |  |  |  |
| Ligand |  |  |  |  |
| R.M.S. deviations |  |  |  |  |
| Bond lengths (Å) | 0.01 |  |  |  |
| Bond angles (°) | 1.06 |  |  |  |
| **Validation** |  |  |  |  |
| MolProbity score |  |  |  |  |
| Clashscore | 14.79 |  |  |  |
| Poor rotamers (%) | 2.07 |  |  |  |
| **Ramachandran plot** |  |  |  |  |
| Favored (%) | 97.38 |  |  |  |
| Allowed (%) | 2.54 |  |  |  |
| Disallowed (%) | 0.08 |  |  |  |

**Table 1 (Cont). Cryo-EM data collection, refinement, and validation statistics.**
|  | Ankyrin-1 complex<br>Class 5 | Ankyrin-1 complex<br>Class 6 | Free band 3 |
| --- | --- | --- | --- |
|  | (EMDB-TBA)<br>(PDB-TBA) | (EMDB-TBA)<br>(PDB-TBA) | (EMDB-TBA)<br>(PDB-TBA) |
| **Data collection and processing** |  |  |  |
| Magnification | 105,000x | 105,000x | 105,000x |
| Voltage (kV) | 300 | 300 | 300 |
| Electron exposure (e-/Å<sup>2</sup>) | 58 | 58 | 58 |
| Defocus range (µm) | -0.5 to -1.5 | -0.5 to -1.5 | -0.5 to -1.5 |
| Pixel size (Å) | 0.83 | 0.83 | 0.83 |
| Symmetry imposed | C1 | C1 | C2 |
| Initial micrographs (no.) | 15,756 | 15,756 | 15,756 |
| Final micrographs (no.) | 14,464 | 14,464 | 14,464 |
| Initial particle images (no.) | 1,781,329 | 1,781,329 | 1,781,329 |
| Final particle images (no.) | 91,366 | 124,899 | 37,066 |
| Map resolution (Å) | 3.2 | 3.1 | 2.8 |
| FSC threshold |  |  |  |
| Map sharpening B factor (Å<sup>2</sup>) | 54.8 | 55.9 | 65.0 |
| **Model composition** |  |  |  |
| Nonhydrogen atoms |  |  | 18,854 |
| Protein residues |  |  | 1,108 |
| Ligands |  |  | 21 |
| **B factors (Å<sup>2</sup>)** |  |  |  |
| Protein |  |  | 52.67 |
| Ligand |  |  |  |
| R.M.S. deviations |  |  |  |
| Bond lengths (Å) |  |  | 0.003 |
| Bond angles (°) |  |  | 0.7 |
| **Validation** |  |  |  |
| MolProbity score |  |  |  |
| Clashscore |  |  | 5.35 |
| Poor rotamers (%) |  |  | 1.16 |
| **Ramachandran plot** |  |  |  |
| Favored (%) |  |  | 98.0 |
| Allowed (%) |  |  | 2.0 |
| Disallowed (%) |  |  | 0.0 |

#### Motion correction, CTF estimation and micrograph curation

Patch-based motion correction and dose-weighting of 14464 cryo-EM movies was carried out in cryoSPARC (v3.2) using the Patch Motion job type. Patch-based CTF estimation was performed on the aligned averages using Patch CTF.

#### Particle picking

An initial round of exploratory processing was performed using Blob Picker. Ranking the micrographs by the number of particles remaining after 2D classification, 58 exemplary micrographs were selected for manual picking in order to create a training dataset for the Topaz neural network-based picker^48^. 1510 particles were manually picked from 58 micrographs. Topaz was run in training mode using a downsampling factor of 6, and an estimated number of particles per micrograph of 200. The resulting model was used to pick particles using Topaz Extract. 1.8M particles were initially extracted in a box of 600 pixels, and Fourier cropped to 150 pixels for initial cleanup (2D and initial 3D classification).

#### 2D classification

Multiple rounds of 2D classification were performed in order to isolate homogeneous subsets of particles to use for ab initio reconstruction of compositionally distinct classes. The 48 most occupied 2D classes from the initial round of 2D classification are presented in **Extended Data Fig. 2**.

#### Initial model generation

Heterogeneous ab initio reconstruction was performed on a variety of subsets of particles selected by 2D classification, in order to generate a diverse range of initial models representing distinct 3D classes in the data for further classification.

#### Initial 3D classification

Heterogeneous refinement was performed using the Topaz picked initial particle stack of 1.8M particles, with 12 representative initial reconstructions as map inputs. 4 well defined, high-resolution classes of the ankyrin complex were identified, as well as several well-defined classes of smaller particles, most of which were identified as free band 3 dimers. 849k particles corresponding to the ankyrin complex were re-extracted with recentering in a 600 px box.

#### Consensus refinement

An initial refinement of all 849k ankyrin complex particles was performed using non-uniform refinement with on-the-fly refinement of per particle defocus, and global refinement of beam tilt and trefoil aberrations, giving an initial consensus refinement with a resolution of 2.6 Å.

#### Sub-classification of ankyrin complex particles

Based on sub-classification of each of the original four classes, six non-redundant classes were identified. The 849k stack of clean ankyrin complex particles from the consensus refinement was subjected to heterogenous refinement using these six classes as inputs, as well as five decoy classes, using an initial lowpass filter of 50 Å and a batch size per class of 5000 particles. The composition and occupancy of the resulting classes is summarized in Figure S5.

#### Sub-classification using 3D Variability Analysis

In the case of both Band 3-I and AQP1, another approach was taken to isolate a set of particles with improved density for the mobile component (**Extended Data Fig. 7**). First, a local refinement with a mask around the mobile component was performed, yielding a relatively poor resolution, noisy local reconstruction (4.5Å in the case of Band 3-I starting from the consensus refinement). Then, 3D variability analysis was performed as implemented in CryoSPARC 3.2, selecting five modes, with a filter resolution of 4.5Å. Reconstructions were calculated along each mode, and a mode corresponding to an order-disorder transition of the mobile sub-region was identified. This mode was then used (using the 3D Variabilty Analysis Display job type) to split the particles into 20 clusters, and a reconstruction calculated for each. Particles belonging to clusters with well-defined, high-resolution features were combined, re-aligned globally by non-uniform refinement, and subjected to another round of local refinement, in the case of Band 3-I yielding a map with a resolution of 3.3 Å.

#### Refinement of ankyrin complex classes

Classes 1, 2 and 3 were individually subjected to high resolution refinement using non-uniform regularization, resulting in reconstructions with resolutions of 3.0, 2.8 and 3.0 Å, respectively.

#### Processing of free band 3 dimer particles

In a separate processing pathway, after identification of the free band 3 dimer class using multi class ab initio reconstruction, several successive rounds of heterogeneous refinement were performed using a single band 3 dimer class and eight random density decoys, against the original 1.8M particle stack. This allowed isolation of a subset of 37k particles which were re-extracted in a 320px box and refined with non-uniform regularization to obtain a reconstruction at 2.8 Å resolution.

#### Local refinements and generation of composite maps

For the consensus refinement and each refined class, local refinements using several different masks were performed to generate local reconstructions with improved density. In order to aid model building of the complex, composite maps were generated by aligning all the local maps to the global reconstruction, and then combining them by taking the maximum value at each voxel using UCSF Chimera^49^.

#### Map post-processing

Maps were sharpened by automated B-factor sharpening, as implemented in cryoSPARC.

#### Atomic model building and refinement

An initial model for protein 4.2 was generated using trROSETTA^50^. Ankyrin-1^51^ (PDB ID: 1N11), Band 3^52,53^ (PDB IDs: 4YZF & 1HYN) and aquaporin crystal structures^54^ (PDB ID: 4CSK) were used as initial models for building the corresponding proteins. Initial models for RhCE and RhAG were generated by threading using CHAINSAW^55^ as implemented in the CCP4 package^56^, using the structure of the RhCG homotrimer as a template^27^ (PDB ID: 3HD6). Initial models were placed in the composite map and fit as rigid bodies using UCSF Chimera^49^. Each model was then manually extended and completed in COOT, followed by refinement using phenix.real_space_refine^57^. Figures were prepared using UCSF ChimeraX^58^. Refinement and validation statistics are provided in **Table 1**.

#### Cryo-ET data processing

Tilt images were frame aligned with MotionCor2^59^ without patches or dose weighting. Frame aligned tilt images were used for fiducial-less tilt-series alignment in Appion-Protomo^60,61,62^. Aligned tilt-series were dose weighted in Appion-Protomo using equation 3 in Grant & Grigorieff^63^ reconstructed for initial visualization with Tomo3D^64,65^ SIRT and separately without dose weighting with Warp^66^. 7 binned by 4 Warp tomograms (pixel size 8.308 Å) were used to train an IsoNet^67^ model (where 300 sub-tomograms were extracted at 963 voxels after masking with z-axis cropping to remove void volumes, then trained over 35 iterations), which was then applied to each tomogram for visualization with IMOD^68^.

#### Cryo-EM data validation

Map resolution was estimated using the gold-standard FSC=0.143 criterion, calculated from half maps using a soft mask in cryoSPARC 3.2. Detailed validation statistics are provided in **Table 1**.

## Acknowledgments

CryoEM data were collected at Columbia Cryo-EM facility and at the Simons Electron Microscopy Center (SEMC), directed by Dr Bridget Carragher and Dr Clint Potter, with the assistance of staff from both SEMC and the Columbia University Cryo-Electron Microscopy Center. Bob Grassucci and Zhening Zhang from the Columbia CryoEM Center assisted with data collection. The authors thank Lorenzo Maso for generating the schematic representation of the ankyrin-1 complex (Fig. 1C) and Dr. Filippo Mancia and his lab members for support and critical reading of the manuscript. We are grateful to Dr. Paola Berto and Dr. Lucia Barazzuol, Department of Biomedical Sciences, University of Padova, for their support in the preparation of the ghost membranes. We thank the “Centro Transfusionale dell’Azienda Ospedaliera Università di Padova” and the individuals who donated their blood from which the ankyrin complex was purified.

## Author contributions

F.V. with the assistance of L.Y.Y. performed protein preparation for structural analysis. F.V. and T.C. prepared membranes from human erythrocytes. F.V. screened and optimized sample vitrification, and generated cryo-EM data. Tomography data analysis was performed by J.D.J. and A.J.N. Single particle cryo-EM data analysis and model building was performed by O.B.C. The manuscript was written by O.B.C. and F.V. with input from T.C. Figures were prepared by K.K and F.V.

## Data and materials availability

The electron density maps and model of the ankyrin-1 complex have been deposited in the Electron Microscopy Data Bank with ID EMD-XXX and in the PDB with XXX, respectively.

---

**Extended Data Fig. 1:**
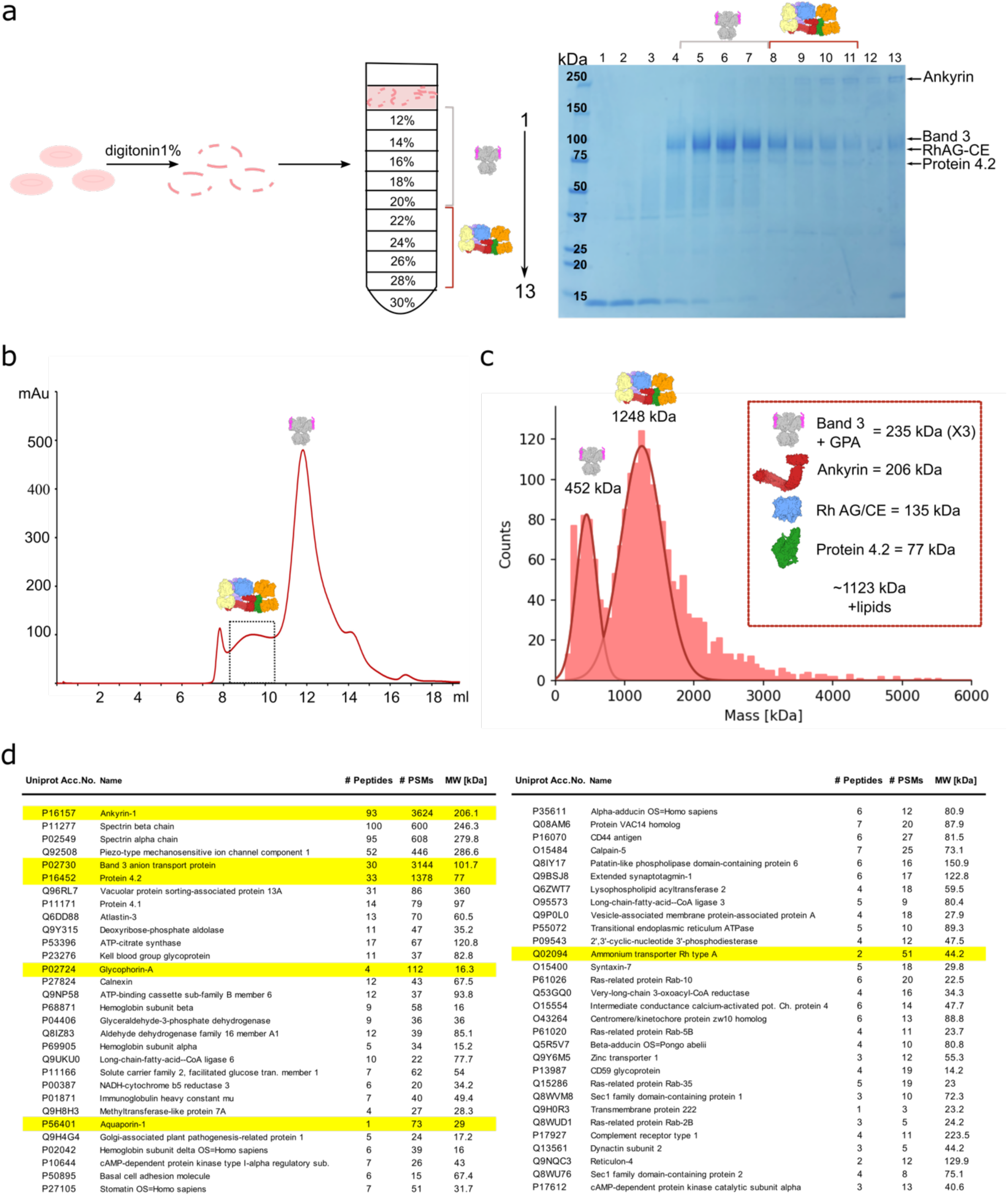
Purification and biochemical characterization of human erythrocyte ankyrin-1 complex. **a,** Human erythrocyte ghost membranes, represented as pink empty erythrocytes, were solubilized using 1% (w/v) digitonin. Membrane fragments were then added in a 12-30% glycerol gradient. The 13 fractions collected were loaded in an SDS-PAGE gel. Gradient fractions containing the complex are marked with a red line and small representations of the complex and the free band 3 both in the gradient and in the gel. **b,** Size exclusion chromatography of the fractions containing the complex show two different peaks, one eluting around 9.5 mL and the other at 12.2 mL. They correspond to the ankyrin-1 complex and the isolated band 3-glyophorin A complexes respectively. The peak fractions of the ankyrin-1 complex were collected and concentrated for cryo-EM sample preparation. **c,** Mass photometry analysis of the sample used for cryo-EM. The main peak represents a population of proteins with a size around 1.2 MDa. The peak at 450 kDa likely represents the band 3-glycophorin A dimer population. In the red square the list of the components of the ankyrin-1 complex is provided with their respective molecular weights. **d,** Mass spectrometry of sample used for structural analysis. MS detected proteins are shown in order of decreasing confidence. All characterized ankyrin-1 complex components are listed (yellow). Most of the contaminating proteins are plasma membrane proteins.

**Extended Data Fig. 2:**
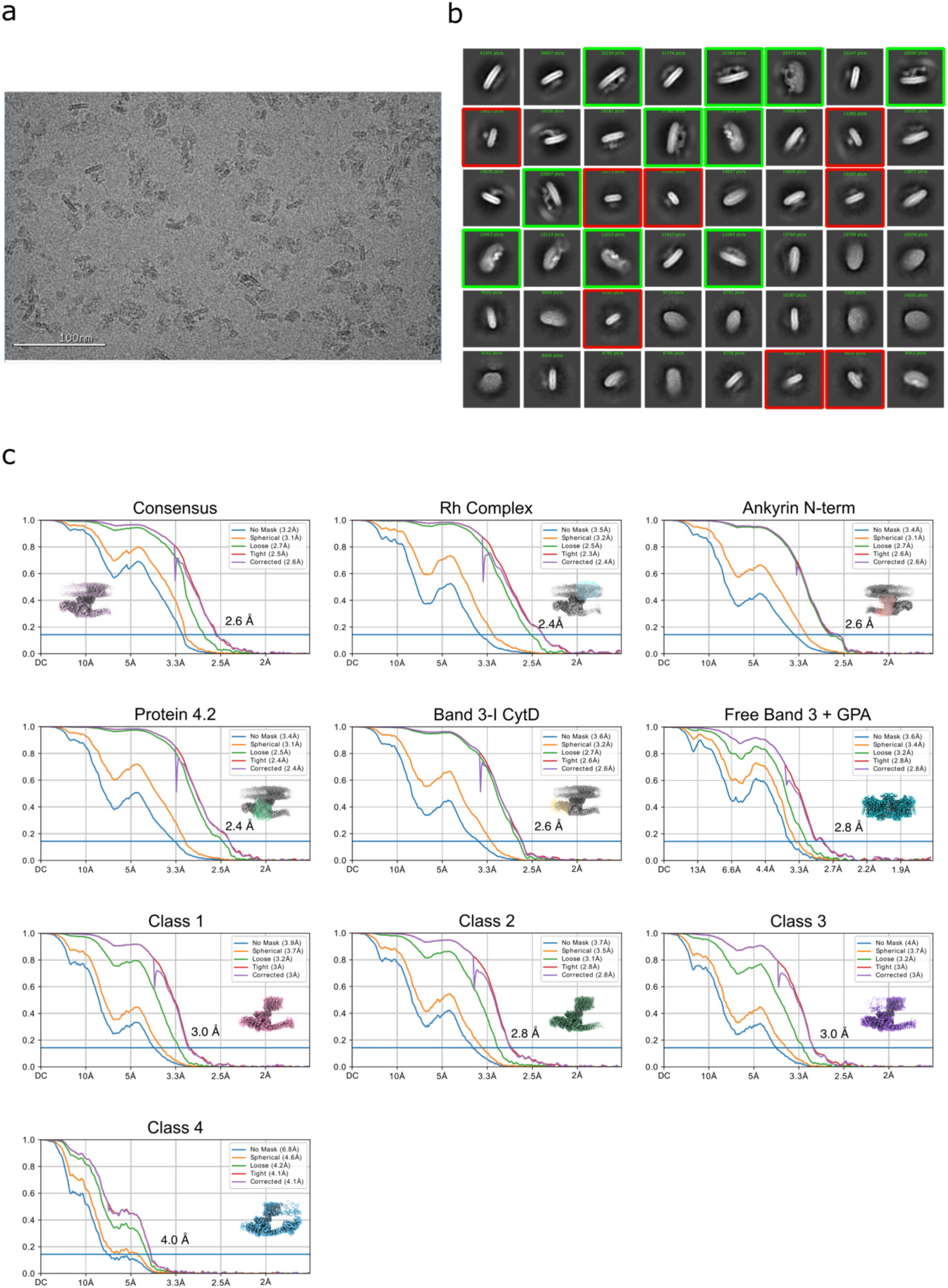
Single particle cryo-EM structure determination. **a,** A representative micrograph. **b,** 2D class average of particles initially picked with Topaz (∼1.8 M), ordered by population. 2D class averages of free band 3 (red box) and ankyrin-1 complex (green box) are highlighted. **c,** Fourier shell correlation (FSC) curves. Maps shown as inset, with refinement mask in the case of local refinements.

**Extended Data Fig. 3:**
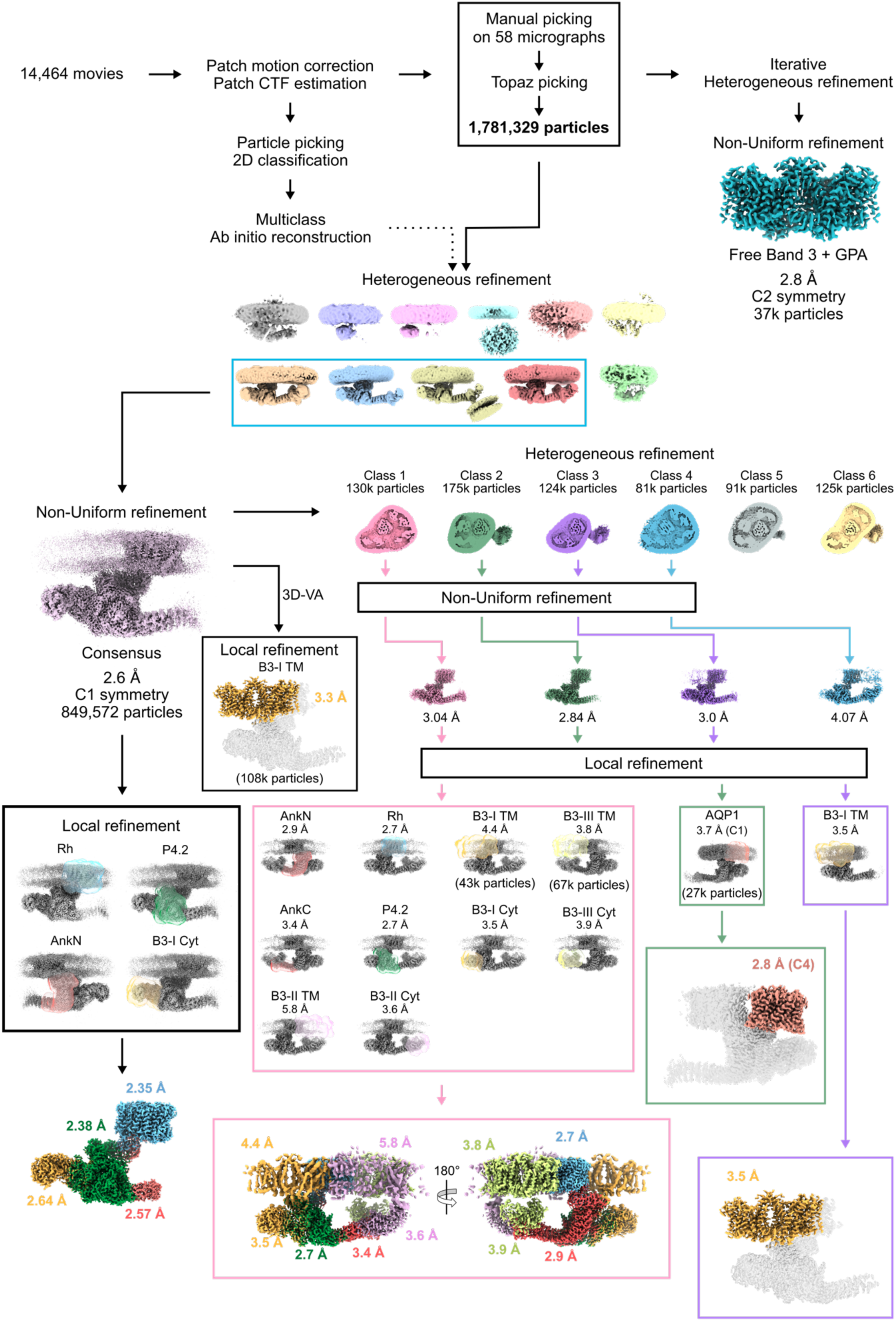
Cryo-EM workflow and analysis of the ankyrin-1 complex. Flowchart outlining cryo-EM image acquisition and processing performed to obtain the structure of ankyrin-1 complex. All processing was performed using CryoSPARC v.3.2 (see Methods for details).

**Extended Data Fig. 4:**
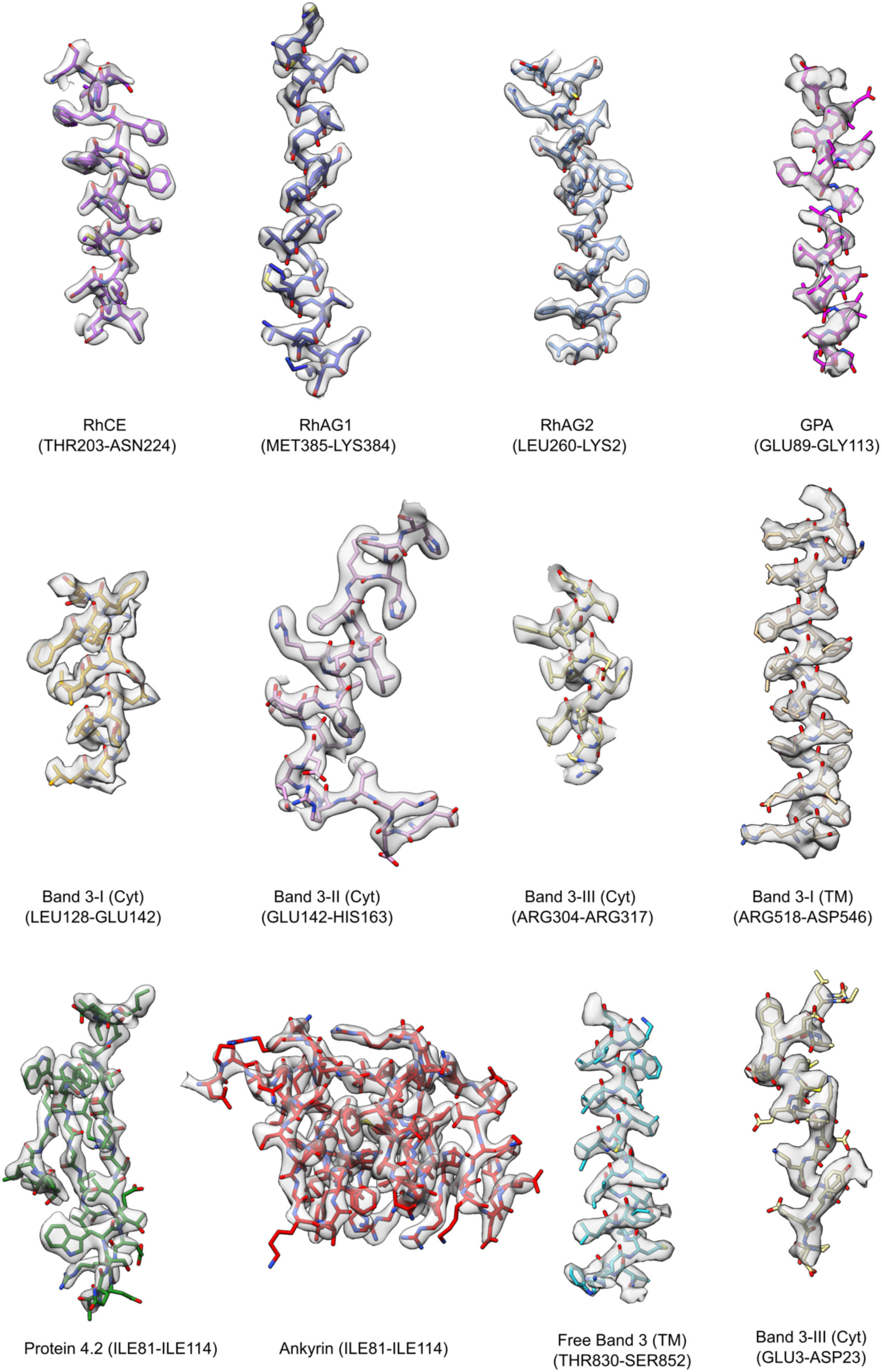
Model/map fit. Cryo-EM densities (transparent gray surface) are shown with corresponding segments of the atomic model; sidechains are rendered in stick representation and colored as in Figure 1. The densities for GPA and free band 3 are derived from the free band3 class. Density for band 3-I TM derives from local refinement of Class 3. For all the other images we used the density from local refinement of the consensus refinement.

**Extended Data Fig. 5:**
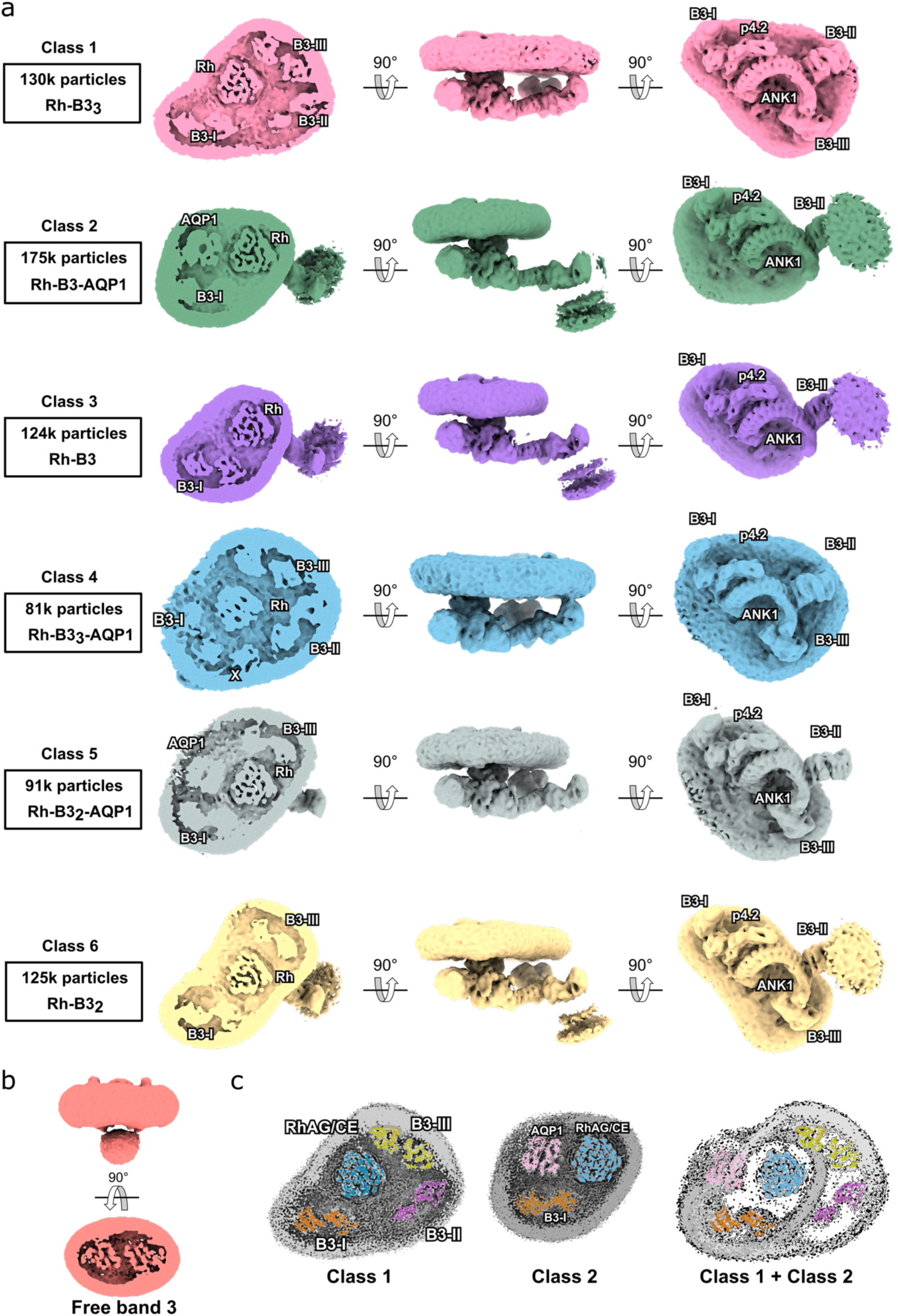
Ankyrin-1 complex classes. **a,** Different views of six main classes of the ankyrin-1 complex. Class 1 contains an (RhAG)_2_(RhCE) heterotrimer, ankyrin-1, protein 4.2, and three band 3 dimers (I, II, III). Class 2 exhibits a smaller micelle, retaining the core (RhAG)_2_(RhCE)(Ank1)(P4.2) architecture, but including only one band 3 dimer (B3-I), bound to protein 4.2, and aquaporin. Class 3 contains an (RhAG)_2_(RhCE) heterotrimer, and a single band 3 (B3-I). Class 4 has a bigger micelle and contains an (RhAG)_2_(RhCE) heterotrimer, ankyrin-1, protein 4.2, three band 3 dimers and an unidentified protein “X”. Class 5 contains an (RhAG)_2_(RhCE) heterotrimer, ankyrin-1, protein 4.2, aquaporin and Band 3-I and Band 3-III. Class 6 contains an (RhAG)_2_(RhCE) heterotrimer, ankyrin-1, protein 4.2, and Band 3-I and Band 3-III. In all the six classes the cytosolic domain of Band 3-II is present. **b,** Free band 3-GPA class shown from the side and from the top. **c,** Comparison of Class 1 & Class 2. Rh is colored in blue, Band 3-I in orange, Band 3-II in lilac, Band 3-III in yellow and aquaporin in pink. Superposition of Class 1 and Class 2 shows that presence of the additional two band 3 dimers does not exclude the presence of aquaporin.

**Extended Data Fig. 6:**
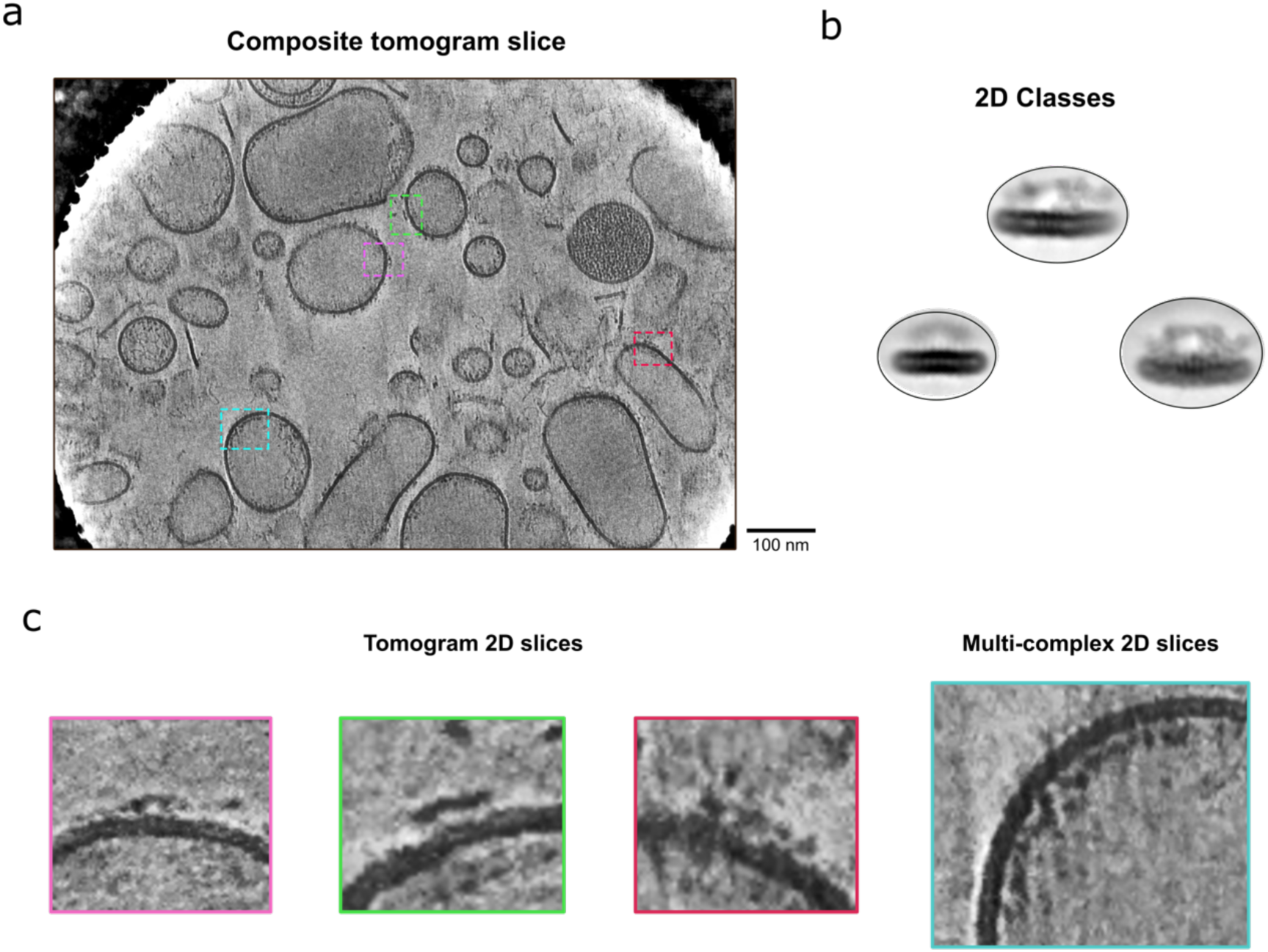
Cryo-ET of native membrane vesicles. **a,** A composite slice from the tomogram shown in Video S1 with several densities resembling ankyrin-1 complex side-views boxed out. **b,** SPA 2D class side-views for comparison with the tomogram slices. **c,** Magnified tomogram slices from the boxed-out regions in A. The multi-complex 2D slice on the right shows one of many visible strings of ankyrin-1 complex-like densities suggesting potential higher order assemblies. Tomogram slices are 8.3 Å thick.

**Extended Data Fig. 7:**
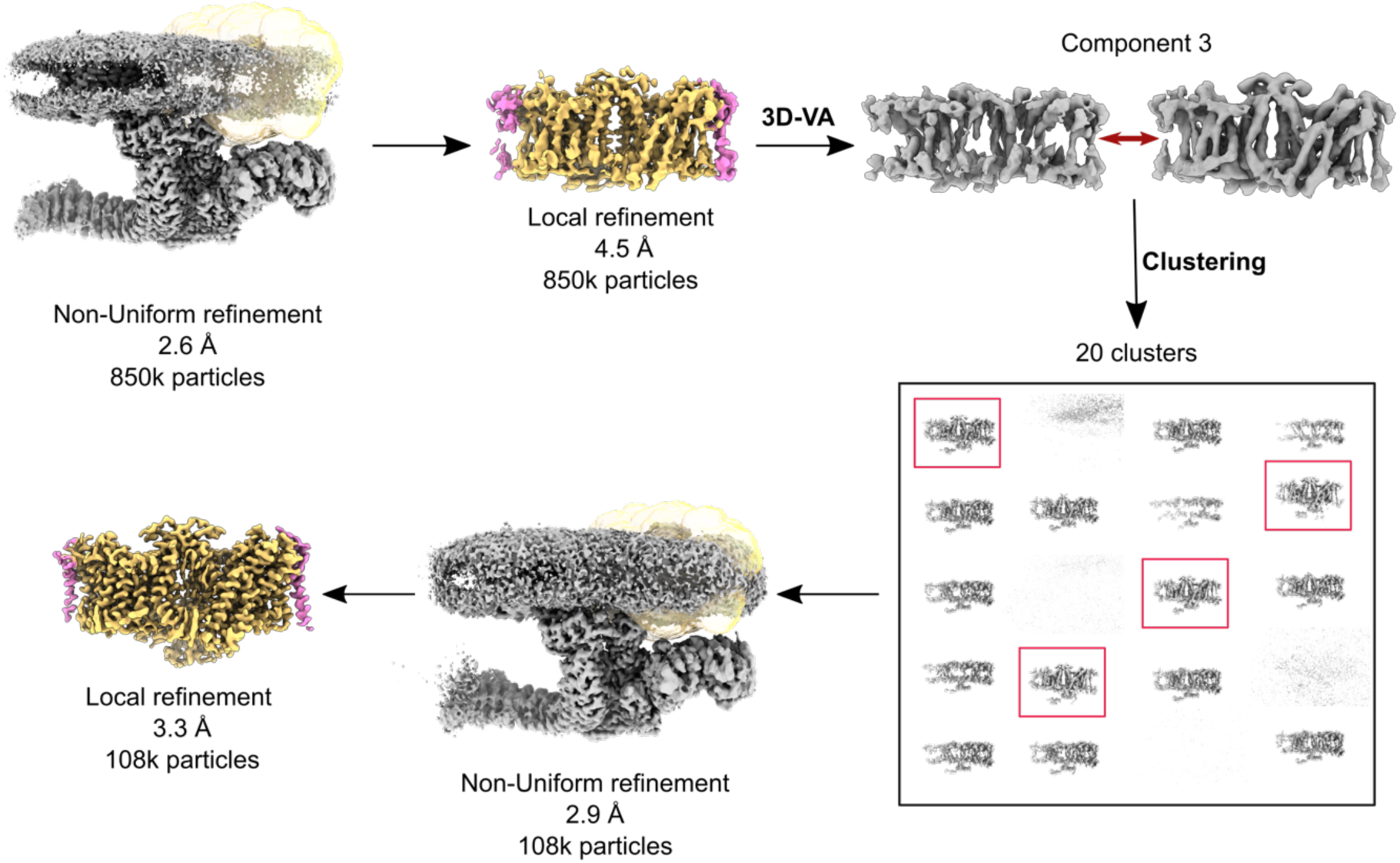
Cryo-EM workflow for sub-classification of Band 3-I from the consensus refinement. Flowchart outlining cryo-EM processing performed to improve the structure of Band 3-I (orange and the bound GPA proteins (magenta). All processing was performed using CryoSPARC v.3.2. A similar procedure was used to improve the density of AQP1 starting from Class 2.

**Extended Data Fig. 8:**
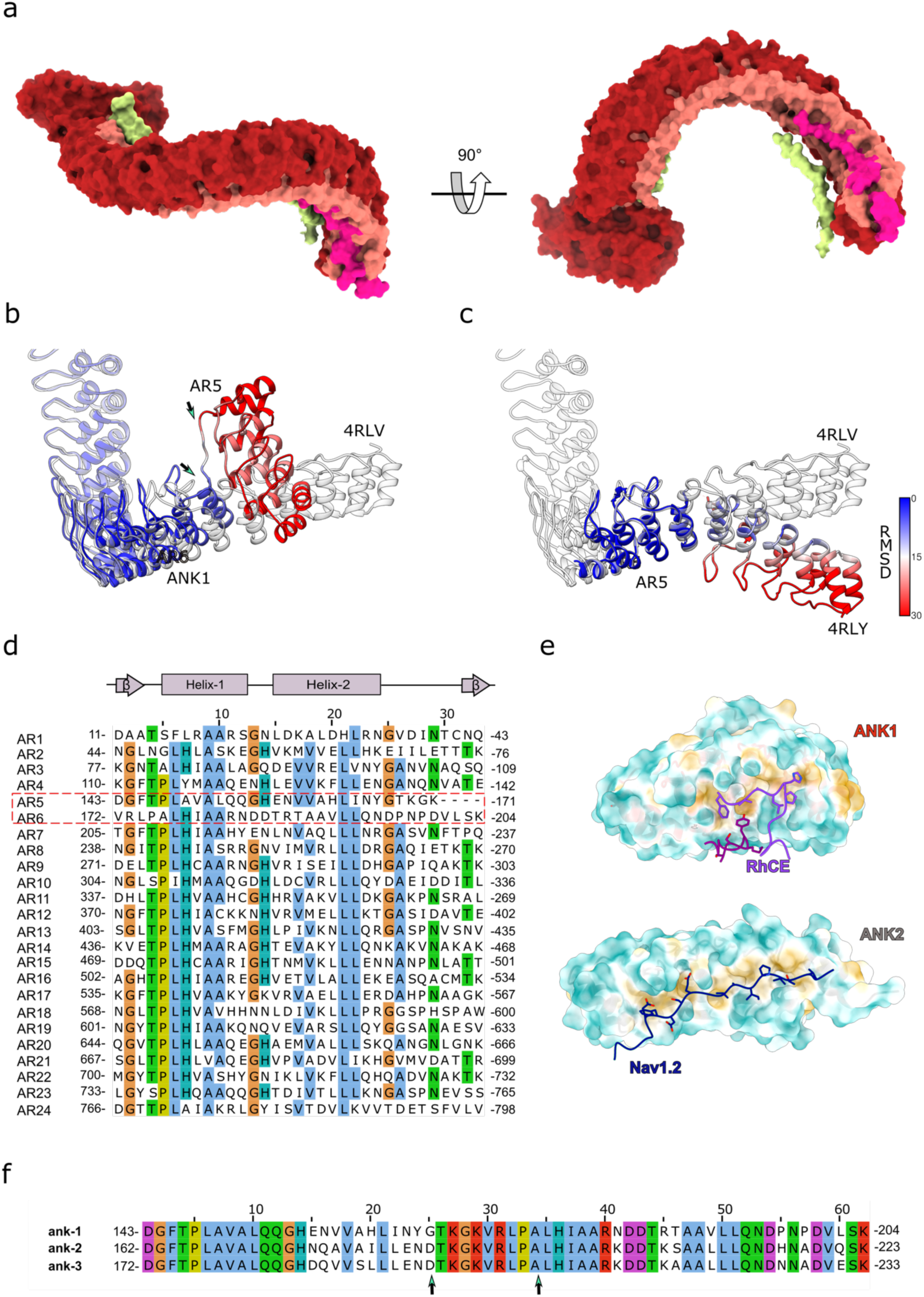
Conformation of membrane-associated ankyrin-1. **a,** Ankyrin structure shown as a molecular surface with the inner groove depicted in pink, the convex outer surface in red, in yellow the N-terminal peptide of Band 3-III and in magenta the C-terminal linker of ankyrin-1. **b,** Structural alignment between ankyrin-1 structure (colored by RMSD) and ankyrin-2 (in gray). The first 5 repeats of Ank1 are dramatically rearranged compared to their position in the Ank2 structure. Arrows indicate AR5 and 6. **c,** Structural alignment between ankyrin-2 (gray) and ankyrin-2 bound to Nav1.2 peptide (colored by RMSD), shows that flexibility at this interface also occurs in ankyrin-2. **d,** Sequence alignment of all the 24 ankyrin repeats of Ank1 using MUSCLE(*46*) and visualized and colored using Jalview with the ClustalX color scheme. The sequences for AR5 and 6 are outlined in red. **e,** Hydrophobic surface calculated for the first 5 ankyrin repeats (AR1-5) of Ank1 (top) and Ank2 (bottom). RhCE N-terminal (magenta) and C-terminal fragments (purple) are displayed in stick representation. In the lower figure Ank2 is bound to Nav1.2 peptide (blue). **f,** Sequence of AR5 and AR6 from Ank1, Ank2 and Ank3, aligned using MUSCLE (*46*) and visualized and colored using Jalview with the ClustalX colour scheme. The three sequences are well conserved in AR5 and 6. Arrows indicate the region that rearranges in Ank1.

**Extended Data Fig. 9:**
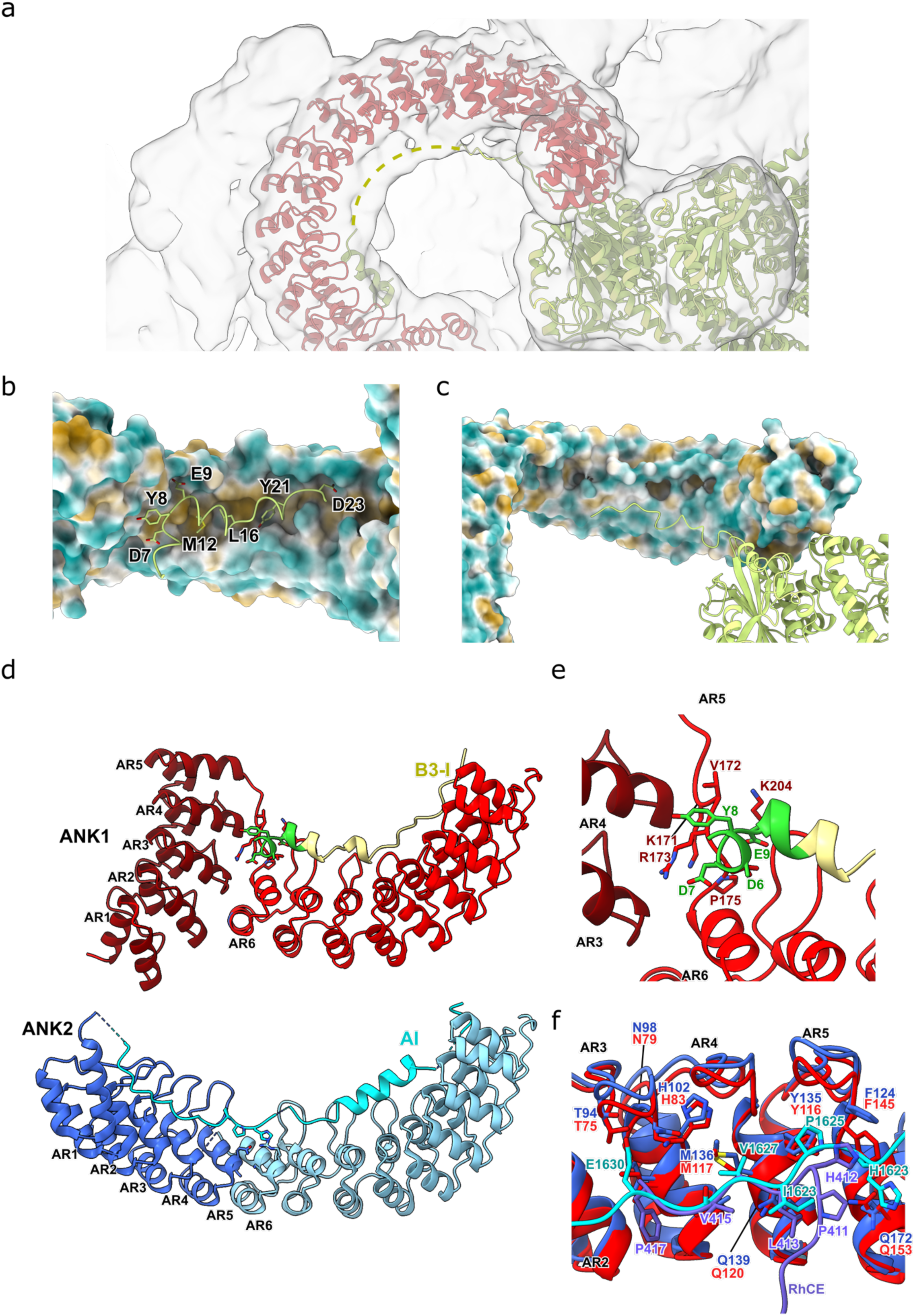
Ankyrin interaction with band 3-III N-terminal peptide. **a**, Map at low density threshold allows tracing of the complete Band 3-III N-terminal peptide (dotted yellow line), that runs back along the inner ankyrin groove. **b-c,** Molecular surface of the ankyrin repeats, colored by hydrophobicity, bound to the N-terminal peptide of Band 3-III (yellow). Residues 2-24 of Band 3-III form an ordered interaction with AR6-10 and the AR5-6 linker. Key residues are displayed as sticks. Band 3-III is shown as yellow ribbons. **d,** Structural comparison between ankyrin-1 (top) and ankyrin-2 structures (PDB: 4Y4D) (bottom). structure of ankyrin-1 (dark red AR1-5 and red AR6-12) is associated with N-terminal peptide of Band 3-I (yellow) and ankyrin-2 (dark blue AR1-6 and light blue AR6-12) is associated with the autoinhibitory domain (AI) of ankyrin-1. **e,** Close up of the AR5-6 of ankyrin-1 in panel S9D. The Band 3-I part colored in green corresponds to the first 11 N-terminal amino acids. **f,** Structural alignment between the first five ankyrin repeats (AR1-5) of ankyrin-1 (dark red) bound to RhCE peptide and ankyrin-2 (dark blue) bound to the autoinhibitory peptide of ankyrin-1. The key conserved residues that mediate the interactions between ANK1 and B3-III and ANK2 and the autoinhibitory region are shown as sticks.

**Extended Data Fig. 10:**
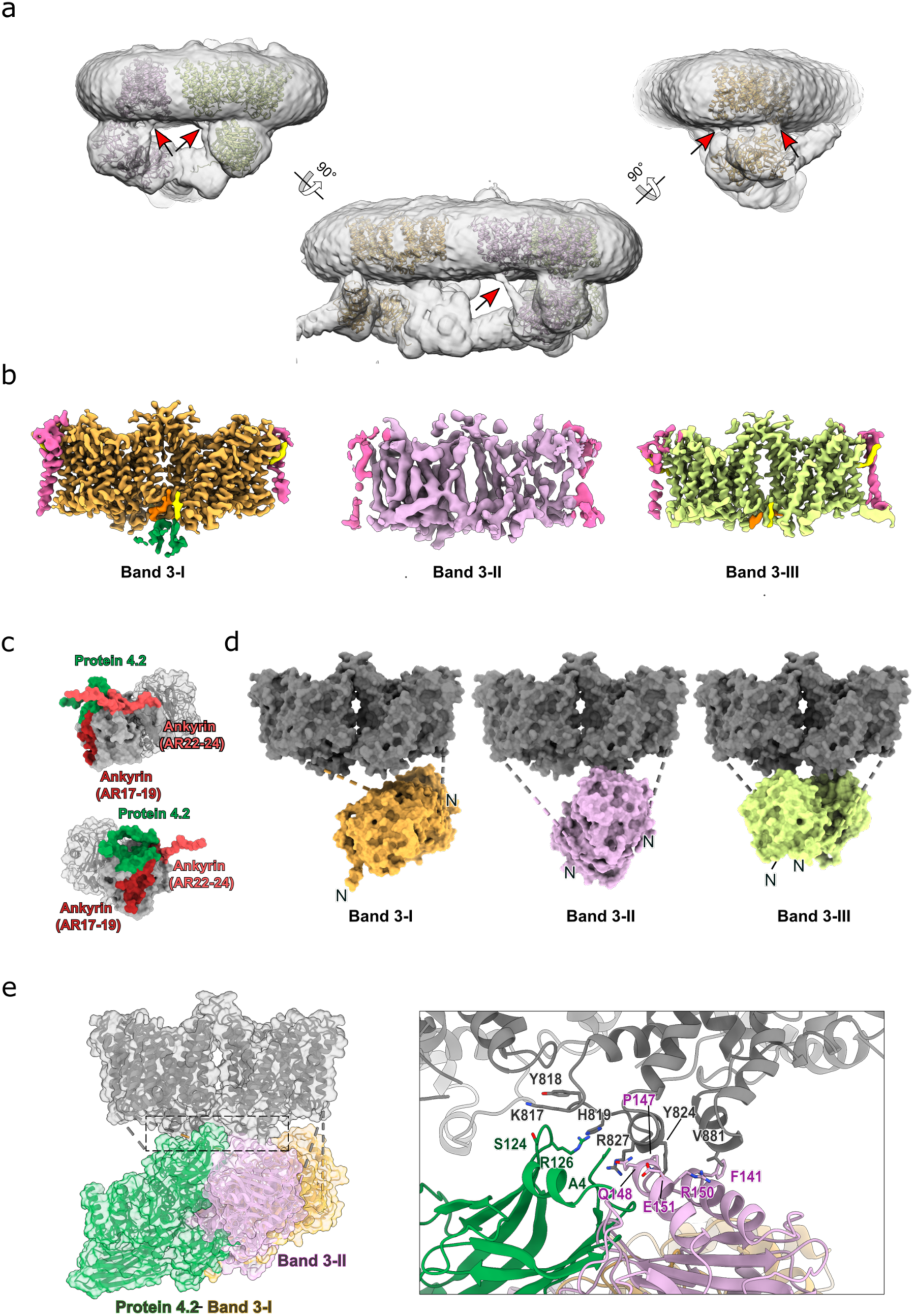
Flexibility of band 3 cytosolic domain. **a,** Class 1 density map at low density threshold allows the identification of band 3 cytoplasmic-transmembrane linkers (marked with red arrows). Models of Band 3 I, II & III are fitted inside the map. **b,** Cryo-EM density map for transmembrane domains of band 3 in ankyrin-1 complex. Focused refinement improved the density of Band 3-I (in Class 3) to 3.5 Å and Band 3-III (in Class1) to 3.8 Å, revealing the presence of one glycophorin A (GPA) monomer bound to each protomer of band 3 (magenta). Even at low resolution (5.8 Å) it was possible to assign the density for Band 3-II and associated GPA monomers. **c,** Two different views of band 3 cytosolic domain represented as molecular surface. Interacting surfaces of band 3 are distinct and non-overlapping. Regions of the band 3 cytosolic domain that interact with proteins in the complex are shown in different colors: green for the region that interact with protein 4.2, red for ankyrin repeats 22-24 and purple for ankyrin repeats 17-19. **d,** Models of the three Band 3 dimers (I, II, III) are displayed as molecular surfaces. The three cytosolic domains of Band 3 proteins in the complex have different orientations. The Band 3-I cytoplasmic domain is inverted with respect to those of Band 3-II and Band 3-III. The TM part of Band 3 is colored gray. **e,** Comparison between Band 3-I (yellow) associated with protein 4.2 (green) and Band 3-II (lilac). The model is displayed as ribbon and the surface is shown in transparency. In the right panel a close view of the interactions between Band 3-II cytosolic domain (lilac) and protein 4.2 (green); the two share the same interaction site with the TM part. Key residues for the interaction are displayed as sticks.

**Extended Data Fig. 11:**
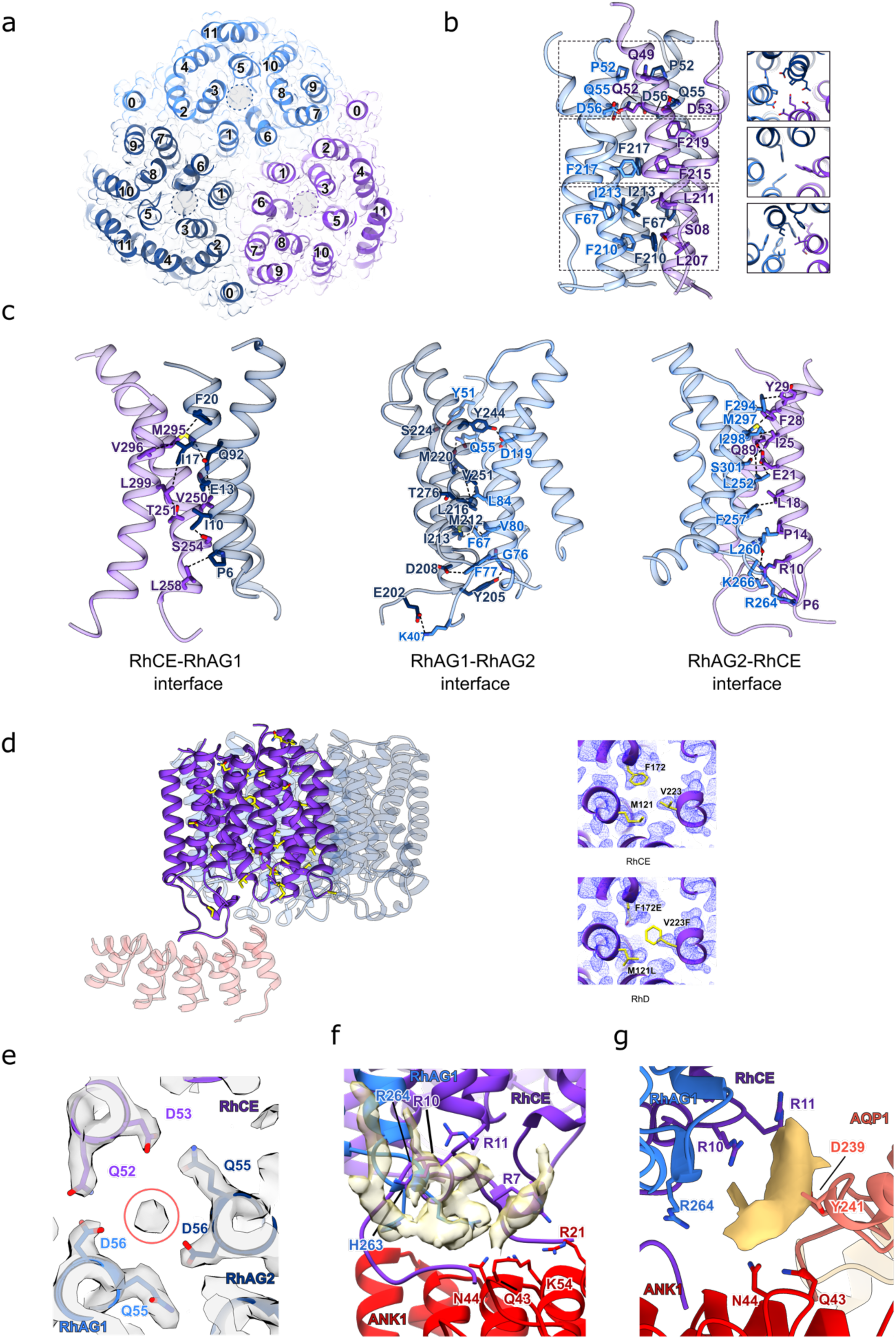
(RhAG)*_2_*(RhCE) trimer interactions. **a,** The structure of (RhAG)_2_(RhCE) trimer as viewed from the cytoplasm, RhCE is colored in purple, RhAG1 in light blue and to RhAG2 in dark blue. The cryo-EM density map of the (RhAG)_2_(RhCE) trimer is shown in transparency. The 12 TM helixes are numbered (0-11) for each Rh molecule. Gray circles shown the pore position in each subunit. **b,** Interface between the three Rh protomers. The key residues that mediate the interactions in the interfaces are shown as sticks. Three different parts of the trimerization interface are show in the right figure viewed from the top. **c,** Interfaces between the three different subunits, with key residues represented as sticks. **d,** Structure of (RhAG)_2_(RhCE) complex and the first five repeats of ankyrin. The amino acids depicted in yellow sticks highlight the sites of variation between RhCE and RhD. Three sites of variation between RhD and RhCE are shown in the right panels, with the density map overlayed. The density map is consistent with the presence of RhCE, not RhD. **e,** Unknown density in the (RhAG)_2_(RhCE) trimer interface coordinated by three aspartates. **f,** Unmodeled lipid densities in a positively charged pocket at the interface of ankyrin-1 (red), RhAG (light blue) and RhCE (purple). Lipid densities are shown as a transparent yellow surface. Potential interacting sidechains are depicted in stick representation. **g,** Unmodeled density at the interface of aquaporin-1 (salmon) with RhCE (purple) and RhAG (light blue). Potential interacting sidechains are depicted in stick representation.

**Movie S1. Cryo-ET of native membrane vesicles.** A composite slice from the tomogram with several densities resembling ankyrin-1 complex side-views boxed out. The multi-complex 2D slice on the right shows one of many visible strings of ankyrin-1 complex-like densities suggesting potential higher order assemblies. Tomogram slices are 8.3 Å thick.

